# A YoeB toxin from *A. tumefaciens* has metal-dependent DNA cleaving activity

**DOI:** 10.1101/795211

**Authors:** Julia McGillick, Jessica R. Ames, Tamiko Murphy, Eswar Reddem, Christina R. Bourne

## Abstract

Toxin-antitoxin (TA) systems, including YoeB-YefM, are important mediators of bacterial physiological changes. *Agrobacterium tumefaciens* YoeB and YefM are similar to that from *E. coli,* and interact as a tight heterotetramer with a K_D_ of 653 pM. We have verified that AtYoeB can perform both ribosome-dependent and –independent RNA cleavage. We have also characterized a newly described metal-dependent and pH-sensitive DNA cleaving ability. We note that this DNA cleaving ability is observed at toxin concentrations as low as 150 nM. The dose-dependence of *in vitro* ribosome-independent RNA and metal-dependent DNA cleavage is equivalent, and requires a ten-fold increase in toxin concentration as opposed to in the presence of the ribosome. The toxin concentration inside bacterial cells is unknown and according to current models, should increase upon activation of YoeB through degradation of the YefM antitoxin. The discovery of general nuclease activity by AtYoeB, and perhaps other YoeB toxins, offers an opportunity to explore the plasticity of this protein fold and its potential role in the evolution of nucleases.

## IMPORTANCE

The importance of these findings is directly related to the *in vivo* concentration of YoeB toxins, which remains unknown. At concentrations greater than 150 nM, the YoeB toxin from *Agrobacterium tumefaciens* mediates both ribosome-independent RNase and metal-dependent DNase activity. While our data indicate the ribosome-independent activities of YoeB toxins are weaker than the ribosome-dependent activity, as the TA system is activated it is expected that the YoeB concentration will be increased, potentially providing a scenario where these ribosome-independent activities may be relevant *in vivo*. Further, these studies highlight the plasticity of this protein fold with respect to general nuclease functionality.

## INTRODUCTION

Toxin-antitoxin (TA) systems are an important mode of intracellular prokaryotic and archaeal regulation.[1-5] They consist of a relatively stable toxin and a more labile antitoxin. The role of the antitoxin is two-fold: firstly, to neutralize the toxin, and, secondly, to serve as a self-repressor for the transcription of its operon.[6, 7] Cellular proteases, such as Lon and ClpP, then degrade the antitoxin under various “stressful” conditions.[6-12] This degradation allows existing toxin to be freed from its cognate antitoxin, leading to alterations in cell physiology as well as increased transcription of the bicistronic TA operon. The increased transcription coupled with antitoxin degradation results in increased availability of intracellular toxin.[8, 13] TA systems are classified into six categories (Types I-VI) based on the mechanism by which the antitoxin neutralizes the toxin.[3, 10] These systems are found encoded on phages,[14] plasmids,[15, 16] and chromosomes.[1] Type II TA systems are the most widely studied and contain both a protein toxin and antitoxin.[4, 17] Type II toxins generally inhibit translation (HipA,[16] VapC,[18] RelE,[19, 20] Doc,[21] HicA,[22] and MazF,[23, 24]), while a few toxins such as Par#,[25] CcdB,[26] and Fic,[27] inhibit the DNA supercoiling gyrase enzyme. Despite the apparent similarity in functions, type II TA systems have proven to be particularly challenging to subdivide due to the relatively low sequence similarity and inconsistencies between toxin and antitoxin annotations.[1, 2, 28-32] For example, the YoeB toxin belongs to the Rel-Par superfamily, while the cognate YefM antitoxin is a member of the Phd superfamily.[33] A large number of type II TA systems are classified as members of the Rel-Par superfamily, which consists of toxins that are both plasmid-derived and chromosomally encoded. This superfamily is of particular interest as members are very structurally similar but are noted to mediate different alterations to cell function.

While plasmid-encoded TA systems have been linked to post-segregational killing [15] and abortive infection,[34] the role of chromosomally-encoded TA systems is a focus of on-going studies. Potential functions for these systems, which we note are not mutually exclusive nor necessarily shared among all TA systems, include mediating responses to specific stresses, altruistic cell death, or protection from invading genetic material.[35-37] Recent work from our lab has shown that the chromosomal ParE toxin from *P. aeruginosa* can exert both protective and fatal effects on the cell.[38] Additionally, the chromosomally-encoded YoeB toxin from *E. coli* has also been well-studied to reveal a dimeric toxin that interacts with the ribosome to mediate mRNA cleavage, thereby altering available transcripts.[39-44] Previous literature on TA systems found in *Agrobacterium tumefaciens* has focused only on tumor-inducing plasmid-borne TA systems rather than those that are chromosomal.[45-48] In this paper, we explore a chromosomally-encoded proposed Par-type toxin from *A. tumefaciens.* The current study demonstrates that this toxin shares sequence and structural similarities, as well as functional activities, with the well-studied *E coli* ribonuclease YoeB toxin. Through these studies we discovered a previously uncharacterized general nuclease activity imparting the *in vitro* ability to degrade DNA in addition to RNA. It is feasible that additional YoeB toxins from other bacteria share this catalytic ability and suggests an evolutionary path for this toxin family that includes DNA degradation.

## MATERIALS AND METHODS

Protein sequence alignments were carried out using UCSF Chimera, and prepared as a figure using ESpript 3.[49] All protein structure figures and analysis of contacts were made using UCSF Chimera.[50]

### Cloning, Expression, and Purification

The AtYoeB and AtYefM genes were amplified from genomic DNA. The YoeB toxin was cloned into a modified pET-28(a) vector containing a C-terminal GST fusion affinity tag in addition to a 6× His tag, while the YefM antitoxin was cloned into the pET15b vector. These constructs were also cloned into the pET-Duet vector, with the AtYefM protein placed in the first multiple cloning site with an N-terminal 6× His tag, \while the second multiple cloning site contained untagged AtYoeB. Each construct was verified by Sanger sequencing (see **Table S1** for sequences, strains, and plasmids used).

These expression clones were transformed into BL21 DE3 *E. coli* and propagated in Lysogeny broth (Miller, Difco) at 37 °C, 200 rpm until the OD_600_ measured ∼0.6. The temperature was then dropped to 16 °C, and protein expression was induced with the addition of 0.5 to 1 mM isopropyl β-D-1-thiogalactopyranoside (IPTG). The induction extended overnight for AtYoeB and the AtYefM-YoeB co-expression, and 4 to 6 hrs for AtYefM antitoxin. Harvested cultures were resuspended in 50 mM Tris pH 8.5, 300 mM NaCl and mechanically lysed using an EmulsiFlex-C3 (Avestin). Clarified lysate was loaded onto a Roche HisTrap NiNTA column equilibrated in the same buffer, and the His-tagged protein was eluted using imidazole. Relevant fractions of AtYoeB, or AtYefM were desalted into 50 mM Tris pH 7.5, 150 mM NaCl using a HiPrep 26/10 desalting column (GE Healthcare). Cleavage of the GST tag fused to AtYoeB was accomplished through an overnight 4°C incubation with 2 U mg^-1^ HRV3c PreScission protease (Sigma), 1 mM DTT, and 1 mM EDTA. Cleavage of the AtYefM 6×His tag affinity tag was achieved with protein fractions desalted into 50 mM Tris pH 7.5, 150 mM NaCl and incubated overnight at 4°C following the addition of 2 U mg^-1^ of thrombin (Sigma) and 2 mM CaCl_2_.

Cleaved AtYefM was separated from residual uncleaved protein *via* a second purification over NiNTA resin, while cleaved AtYoeB was separated from the larger GST tag by size exclusion chromatography. The final purification step for each sample, including the co-expressed AtYefM-YoeB, utilized a Superdex 75 10/300 GL column (GE Healthcare) equilibrated in 50 mM Tris pH 8, 150 mM NaCl; protein purity was assessed by electrophoresis using 12% or 18% tris-tricine gels. Western blots were carried out using a semi-dry transfer method, blocking with 1% milk in tris-buffered saline (TBS), and washing steps utilizing TBS plus 0.1% Tween-20. Primary antibodies were an anti-pentaHis mouse antibody (Qiagen) and an anti-Strep mouse antibody (IBA), while the secondary antibody as a fluorescently labeled goat anti-mouse antibody (BioRad). Blots were imaged on using a ChemiDoc (BioRad).

### Crystallization of AtYoeB

Crystallization screening was carried out at 293 K in a 96-well plate using sitting drop vapor diffusion crystallography with the aid of a Mosquito crystallization robot (TPP LabTech). The JSCG+, PACT 1, and PACT 2 crystallization screens (Molecular Dimensions) were first tested. Small, intergrown needle-like crystals appeared after 16-18 hours from several conditions with polyethylene glycol (PEG) 3350 as a precipitant. Unfortunately, further optimization by altering the pH, salt and precipitant concentration doesn’t improve shape and size of the crystals. An XP additive screen (MiTeGen) formulated with Anderson-Evans polyoxotungstate (TEW) yielded X-ray diffraction quality crystals obtained after 48 hrs with a maximum size of 150×150×75 μm in a condition of 5% v/v 2-Methyl-2,4-pentanediol (MPD), 10% PEG 6,000, 100 mM 4-(2-hydroxyethyl)-1-piperazineethanesulfonic acid (HEPES) pH 7.5, and 1 mM TEW.

Prior to data collection, AtYoeB crystals were briefly rinsed for about ∼15-30 seconds in cryoprotectant solution supplement with 30 % MPD in the mother liquor and plunged in liquid nitrogen. X-ray diffraction data (extending to 1.75 Å) were collected at 100 K at the SSRL ID14-1 beamline (Stanford, USA) with using an Eiger X 16M detector (Dextris AG). Diffraction data were processed with the program XDS [51] and scaled using AIMLESS [52] from the CCP4 software suite.[53] Molecular replacement was performed with PHASER [54] using PDB entry 2A6Q [42] as a search model. Manual rebuilding of the structure using COOT [55] was alternated with refinement using Phenix.[56] TEW was included after inspection of the initial electron density maps during the final stage of model building and refinement. Water molecules were added following standard criteria based on electron density and proper hydrogen bond interaction geometries. Details of data collection and processing are presented in **Table 1.**

### Differential Scanning Fluorimetry Assay

Purified protein samples in 50 mM Tris pH 8, 150 mM NaCl were diluted to 36 μM and an equal volume mixed with 5 ×SYPRO orange (Invitrogen) to yield 20 μL reactions. Assays were carried out in white 96-well PCR plates using a Roche LightCycler 480II set to the minimum ramp rate from 20 to 95°C with 1 measurement per degree. Resulting melt curves were transformed to the first derivative (d/dT) using manufacturer’s software (StepOne v2.1), visually inspected to reveal the maximum fluorescence change, and replotted using GraphPad Prism (v6.0d). Measurements were repeated three times with essentially identical results, and omission of protein from the samples yielded no change in fluorescence.

### Multi-angle Light Scattering (MALS) Assay

Purified samples of AtYoeB, AtYefM, and co-expressed AtYeoB-YefM were analyzed for absolute molecular weight determination using a Wyatt miniDAWN Treos system integrated with size exclusion chromatography and a refractive index detector maintained at room temperature. The sizing column, a Superdex 200 Increase 10/30 GL (GE Healthcare), was equilibrated in 25 mM Hepes pH 7.5, 150 mM NaCl buffer. The resulting light scattering profiles were analyzed using the ASTRA software (v 6.1, Wyatt Technologies) following manufacturer’s recommendations; concentrations were determined based on the signal from the refractive index detector, as AtYefM contains no tryptophan amino acids, making concentration calculations from absorbance at 280nm error prone. The resulting data were ported to GraphPad Prism (v 6.0d) and replotted for graphic presentation.

### Biolayer Interferometry (BLI) Assay

In order to measure the binding between the toxin and antitoxin, an NiNTA pin (ForteBio) was incubated with 125nM His-AtYefM, followed by titration of concentrations of AtYoeB (after removal of the 6× His tag). All solutions were prepared in a 1× block buffer consisting of 0.5 %BSA, 0.05 %Tween-20 in 50 mM Tris-HCL pH 8.5 and 300 mM NaCl. Controls were included with every measurement, consisting of loading a non-specific protein (His-dihydrofolate reductase) as well as not loading any protein to the NiNTA pin; neither showed appreciable signals indicating interaction with AtYoeB, and the “empty” pin signal was used to correct for the baseline. Pins were regenerated before and between runs by incubations in 10 mM glycine pH 1.7 alternated with 1× block buffer for three cycles of five seconds, followed by recharging in 10 mM NiCl_2_ for 60 seconds. Data were processed with the ForteBio Octet Data Analysis software using best practices. Sufficiently good fits were obtained using a model for a 1:1 stoichiometry (see Fig. S3 for individual assays and calculations).

### DNase Assays

Reactions to measure the ability of a toxin to cleave DNA were assembled by combining equal volumes of 500 ng supercoiled plasmid pBR322 (prepared with Zyppy miniprep kit), 2 mM MgCl_2_, and water with 2× concentration of purified AtYoeB (in 50 mM Tris pH 8, 150 mM NaCl), resulting in 20 μL assay volumes. Reactions were incubated at 37 °C for 30 minutes, mixed with SDS to 1%, and then incubated at 50 °C for 15 minutes. Samples were then mixed with loading dye, applied to a 1% agarose TAE gel containing SYBR safe, and electrophoresed at 80 V for 30 min. To test pH, substrate, or metal dependence, a final concentration of 5 or 10 μM toxin, as noted in the individual experiments, was added to the reaction, and the relevant solutions were varied appropriately. In the case of the pH dependence assay, Bis-Tris buffer at pH 5 or 6 or Tris-HCl at pH 7, 8 or 9 was added to a final concentration of 100 mM. For analysis of topology preferences of DNase activity, substrates included supercoiled pBR322 as listed above, linearized pBR322, or the single-stranded M13 genome (NEB). In the case of the metal dependence assay, MgCl_2_, MnCl_2_, CaCl_2_, or ZnCl_2_ were added to a final concentration of 2 mM; control reactions lacking toxin protein ensured these metal solutions did not contain residual DNA cleaving contaminants.

### RNA Synthesis

RNA was synthesized from SmaI (NEB) linearized plasmid containing the Firefly luciferase gene under control of a T7 promoter (Promega). Linearized DNA was recovered through a standard phenol:chloroform extraction, followed by a back extraction. Glycogen at 10 μg/mL was used as a carrier during an overnight ethanol precipitation at -20 °C as described.(64) Purified DNA substrate was added to the HiScribe™ T7 High Yield RNA Synthesis kit (NEB), and RNA synthesis was carried out according to manufacturer’s directions. The resulting product was electrophoresed on a 1.2% FlashGel™ RNA cassette (Lonza) at 275 V for 8-10 minutes to assess purity. RNA was purified via phenol:chloroform extraction and back extraction following DNase I (NEB) treatment. The final product was recovered via ethanol precipitation.

### Gel-based RNase Assay in the absence of ribosomes

The ability of AtYoeB to cleave RNA in the absence of the ribosome was measured using a gel-based assay. Reactions contained a final concentration of 80 ng Firefly Luciferase (Fluc) RNA, AtYoeB at 0.625 μM to 10 μM in 50 mM Hepes pH 7.5 to 8.5, 150 mM NaCl, and for measurements of Mg^2+^ dependence included DNase I buffer (NEB, used at final concentrations of 10 mM Tris-HCl pH 7.6, 2.5 mM MgCl_2_, 0.5 mM CaCl_2_) or EDTA at 5 mM to chelate metal cations. Control samples contained Fluc RNA with buffer alone, buffer and MgCl_2_, and buffer and EDTA. Reactions were incubated at 37 °C for 30 minutes, after which, the reactions were mixed with formaldehyde loading dye (Lonza). Samples were then loaded onto a 1.2% FlashGel™ RNA cassette (Lonza) and electrophoresed at 275 V for 8 minutes using the Lonza FlashGel™ kit. Product bands were visualized with the manufacturer provided software and quantified with ImageJ.[57]

### Ribosome Dependent Nuclease Assay

We measured the activity of AtYoeB in the presence of the ribosome by monitoring the production of Green Fluorescent Protein (GFP) using the PURExpress In Vitro Protein Synthesis Kit (NEB). Starting substrates were derived from a clone of GFP in a pET28(a) vector (pET28:GFP was a gift from Matthew Bennett, Addgene plasmid # 60733; http://n2t.net/addgene:60733; RRID:Addgene_60733), and reactions contained a final concentration of either 300 ng of linearized GFP DNA or 7.5 μg of GFP RNA (synthesized as described above). Fluorescence was measured every 15 minutes for 2 hours using an excitation wavelength of 485 nm and an emission wavelength of 528 nm in a Synergy H1 Hybrid Multi-Mode Microplate Reader (BioTek). Data were analyzed by subtracting the background fluorescence arising from a reaction with no DNA or RNA substrate to give a corrected fluorescence. The corrected fluorescence at the 2-hour time point was divided by the corrected fluorescence of the positive control (containing only the starting substrate with buffer and no toxin), converted to percentages, and graphed as a function of toxin concentration.

## RESULTS AND DISCUSSION

### Based on sequence and structure, *Atu2017* encodes a YoeB toxin

The genes *Atu2017* (UniProt ID A9CID9) and *Atu2018* (UniProt ID Q7CY23) from *Agrobacterium tumefaciens* strC58 were first identified as a potential TA pair by the Rapid Automated Scan for Toxins and Antitoxins in Bacteria (RASTA-Bacteria).[58] The toxin-antitoxin database (TADB 2.0) predicted *Atu2017* and *Atu2018* to be members of the Par/Rel toxin superfamily fold and the PhD antitoxin superfamily fold.[59] These predictions support an expected YoeB toxin, as the cognate YefM antitoxin is housed within the PhD fold.[60]

The structure of the YoeB toxin from *A. tumefaciens*, herein referred to as AtYoeB, was determined at 1.75 Å resolution (see **Table 1**). YoeB toxins conform to a previously characterized RNase fold consisting of two long helices packed against a twisted four to five-stranded antiparallel beta sheet.[42, 43] When compared to the EcYoeB toxin, there is an overall RMSD of 0.6 Å for the core residues with 1.2 Å deviation overall (**Fig. 1A**). As can be deduced from the protein sequence alignment in Figure 1, these two toxins are 57% identical (76% similar) at the amino acid level, including absolute conservation of the identified EcYoeB catalytic residues noted in previous studies (**Fig. 1**).[41, 43] Similar residues also include C-terminal aromatic amino acids that likely help to stabilize the substrate, such as Phe86 or Tyr88 in AtYoeB, Tyr84 in EcYoeB, and Phe91 in the closely related EcYafQ toxin.[61] Of the 35 different amino acids between EcYoeB and AtYoeB, nine are located at the antitoxin interaction surface. The AtYoeB sequence contains four inserted amino acids, located in a loop between beta-strands 4 and 5 (**Fig. 1A**). This loop was disordered in one of the two copies of AtYoeB present in the crystallographic asymmetric unit; in addition, this loop makes interactions with the tungstate-terillium compound that was critical for bridging important crystal contacts and that was required to obtain usable diffraction quality crystals (**Fig. 1A**).

**Figure 1.**
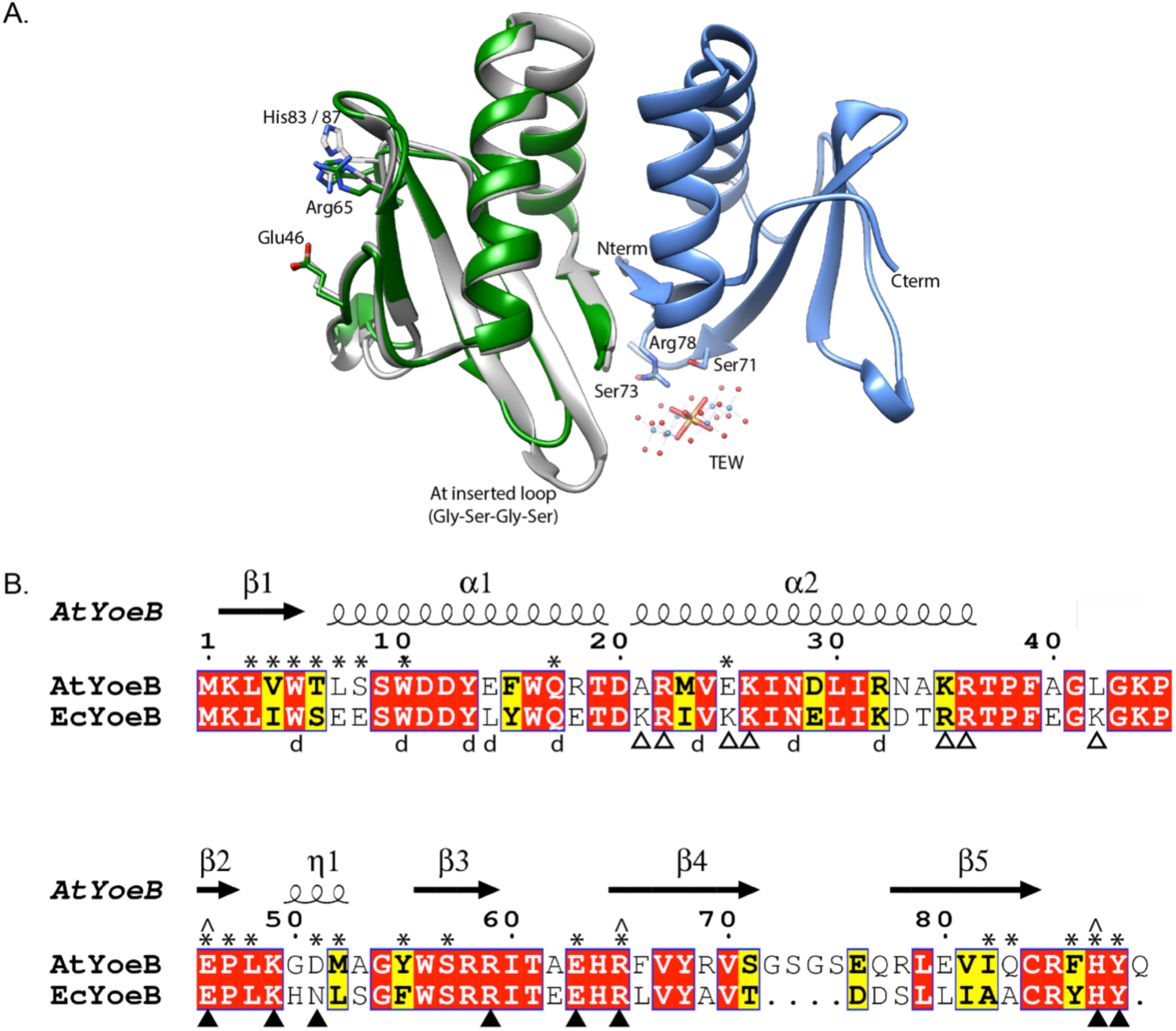
AtYoeB is similar to EcYoeB both in structure and in sequence. **A.** An AtYoeB dimer (blue and grey ribbons) is found in the asymmetric unit of the crystal, with important crystal packing contacts mediated by a tungstate-terillium compound (TEW). The catalytic residues Glu46, Arg65, and a C-terminal His are conserved (shown in stick form, EcYoeB shown as green ribbon). **B.** The sequence of AtYoeB and EcYoeB are 76% similar, depicted with conserved residues in red boxes and similar in yellow boxes. Secondary structure is depicted above the sequence. Asterisks denote amino acids contacting the YefM antitoxin; “d” denotes amino acids at the YoeB dimer interface. Closed triangles denote contacts with the ribosome needed for catalysis, while open triangles mediate weak electrostatic contacts outside of the catalytic region (PDB IDs 6OXA, 6OTR, 6OXI).[2] Antitoxin contacts deduced from superposition of the AtYoeB structure (PDB ID 6N90) onto the *E. coli* YoeB-YefM complex (PDB ID 2A6Q).[3]

The regions of basic charge are maintained, and when superposed with the YoeB structure in the ribosome site the amino acids interacting with the ribosome are also largely conserved.

Rather unusually, YoeB toxins are found as dimers even within the ribosome site, while other ribosome-dependent RNases bind as a monomer, raising questions about the functionality of the second YoeB molecule. In AtYoeB, three of the seven interacting residues in the second molecule are changed from that found in EcYoeB: (Ec to At) Lys21Ala, Lys25Glu, and Lys42Leu (**Fig. 1B**, open triangles). As previously noted, these sites on the second YoeB molecule interact only weakly with the ribosome.[43]

| **6N90** |  |
| --- | --- |
| **Data Collection** |  |
| Resolution Range (Å) | 38.9 – 1.755 (1.81 – 1.755) |
| Space Group | P 1 2<sub>1</sub> 1 |
| Unit Cell | $a=30.9$ Å $b=77.7$ Å $c=36.4$ Å<br>$\beta=92.9^\circ$ |
| Total Reflections | 16748 |
| Completeness (%) | 97.3 (98) |
| I / sigma (I) | 2.61 (1.3) |
| Wilson B-factor | 23.9 |
| R-merge | 0.08 |
| **Refinement** |  |
| R-factor (%) | 21.8 (26.5) |
| R-free (%) | 24.6 (34.1) |
| Number of Atoms | 1591 |
| **Model Quality** |  |
| Number of amino acids,<br>AtYoeB mol 1 / B factor (Å<sup>2</sup>) | 88 / 27.8 |
| Number of amino acids,<br>AtYoeB mol 2 / B factor (Å<sup>2</sup>) | 88 / 28.4 |
| Number of water mols /<br>factor (Å<sup>2</sup>) | 123 / 28.3 |
| Number of TEW /<br>B factor (Å<sup>2</sup>) | 1 / 29.7 |
| **RMS Deviations** |  |
| Bonds (Å) | 0.01 |
| Angles (°) | 1.186 |
| **Ramachandran plot** |  |
| Favored (%) | 93.3 |
| Allowed (%) | 6.7 |
| Coordinate error | 0.24 |
| Clashscore | 2.80 |

Each of these differences results in a loss or reversal of charged interaction. It is expected that differences may arise at the toxin-antitoxin interface (**Fig.** 1B, asterisks) as has been previously observed for the cognate pairing of TA proteins.[62] Given that only one of these amino acids is in contact with the antitoxin, it suggests that these amino acids changes are not driven by the inherent specificity for interacting with their different cognate antitoxins, but rather, that they are random or potentially involved with species-specific toxin functionality.

The YoeB dimer interface is well conserved among this class of toxin, largely solidified by two tryptophan residues from each monomer (conserved at positions 5 and 10 for both At and EcYoeB) forming an “aromatic ring cluster,” in addition to polar interactions mediated by conserved Tyr13, Gln17 and Asn18 side chains (**Fig. S1**). Additional interactions are noted for AtYoeB at positions that differ from its Ec counterpart: (Ec to At) Leu14Glu, Glu18Arg, and Lys32Arg (**Fig. S1**). These changes generate additional strong polar interactions for the AtYoeB dimer.

### The AtYoeB toxin can be expressed in the absence of its cognate antitoxin

The YoeB toxin encoded in *Escherichia coli* (EcYoeB) is noted to be toxic, required co-expression with the cognate antitoxin and purification utilizing denaturation and refolding.[39, 40, 42-44, 62] Interestingly, AtYoeB appears much less toxic during over-expression. We have previously measured the toxicity of AtYoeB to *E. coli* and noted a delayed impact on colony forming units, with minimal impact on existing cells until more than ten hours of overexpression.[63] AtYoeB in the absence of antitoxin could reproducibly yield 3-5 milligrams per liter of LB culture, and with greater than 7 milligrams per liter of culture when the antitoxin is co-expressed (see Methods section for induction details).

### Dimeric AtYoeB and AtYefM interact strongly to produce a heterotetramer

The AtYoeB dimer was found to have a relatively stable melting transition (Tm) value by Differential Scanning Fluorimetry (DSF) of ≅53 °C (**Fig. 2A**). In contrast, the YefM antitoxin has no discernable transition in this assay, indicating that it lacks a hydrophobic core that can undergo denaturation. This result is consistent with the YefM extended helical structures seen previously.[42, 64] The complex of AtYoeB-YefM maintains a signal corresponding to toxin denaturation, and in addition gains a species with a stabilized structure with a Tm of 73.2°C.

**Figure 2.**
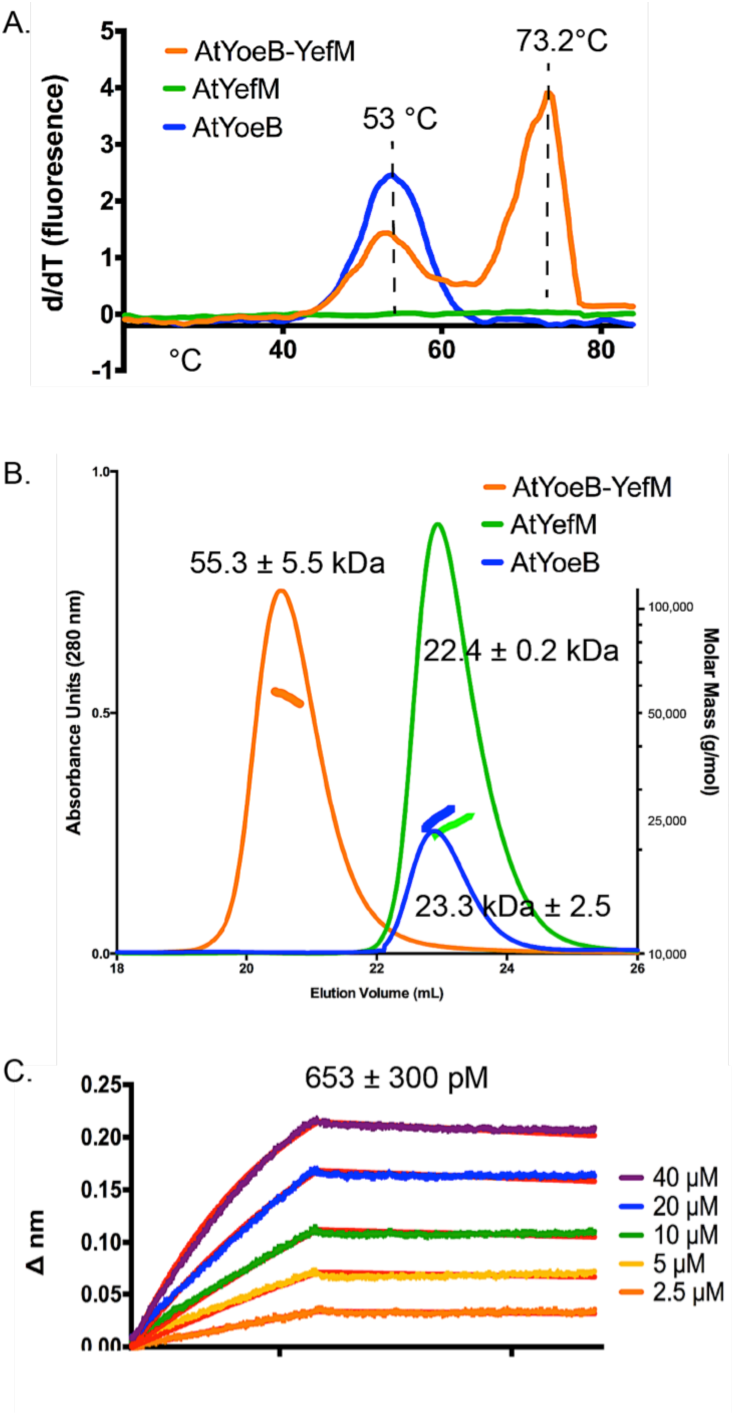
Dimeric AtYoeB interacts with dimeric AtYefM to generate a tightly interacting heterotetramer. **A.** Differential scanning fluorimetry experiments were used to determine that the AtYefM antitoxin (green) does not display an appreciable melting transition in the absence of the AtYoeB toxin (blue), while the complex (orange) yields a single transition with a stabilizing effect versus the toxin alone (*n* = 2). **B.** Size-exclusion multiple angle light scattering (SEC-MALS) establishes the dimeric state of the individual partners and the resulting heterotetramer. AtYefM (green) contained the His affinity tag, while AtYoeB (blue) was analyzed after removal of the GST-His affinity tag (*n* = 2 for each sample). **C.** Biolayer interferometry (BLI) was carried out by capturing His-AtYefM on NiNTA pins and interactions were measured as AtYoeB toxin was titrated (*n* = 3). A one-to-one stoichiometric fit resulted in an interaction strength of 653 pM.

The dimeric state of AtYoeB was verified using MALS, which determined an absolute molecular weight of 23.3 kDa (±2.5, *n*=2) (**Fig. 2B**). While AtYoeB is dimeric, the purified antitoxin was also found to be dimeric by MALS (22.4 ±0.2 kDa, *n*=2) (**Fig. 2B**). To examine the complex stoichiometry, we turned to a co-expression model. Purification relied on the affinity tag of the antitoxin and yielded a single species by size exclusion (**Fig. S3**). This species contains the complex of AtYoeB and AtYefM, and by MALS analysis this complex is a 1:1 interaction resulting in a heterotetramer (55.3 ±5.5 kDa, *n*=3, **Fig. 2B**).

The dimeric AtYoeB toxin interacts strongly with the cognate AtYefM antitoxin, measured by Biolayer Interferometry, yielding a calculated KD of 653 ± 300 pM using a model for a 1:1 fit (**Fig. 2C, S2**). Consistent with this tight interaction, the Tm for this complex by DSF shifts to 73.2 °C (**Fig. 2A**).

### DNase Activity is pH- and metal-dependent but substrate-independent

As part of these studies, we noted AtYoeB-dependent DNA degradation (**Fig. 3**). Previous studies identified the catalytic residues for YoeB toxin degradation of RNA as Glu46, acting as a general base, and the C-terminal His residue (83 in Ec, 87 in At) acting as a general acid. This mechanism indicates that the histidine must be protonated, imparting sensitivity to pH for the reaction. We considered if this mechanism would also be utilized for the degradation of DNA. Consistent with this expectation, there was a sensitivity of DNA degradation to pH. However, the maximum DNA degradation was noted around pH 8-9 (**Fig. 3 and S5**, which includes additional gel images and calculations). At pH 9, potent DNA degrading activity is noted, resulting in only 12% of the DNA remaining supercoiled. At neutral pH 7, an average of around 30% of the DNA is supercoiled, while prominent bands for both nicked and linear DNA substrate are visible and increase as the pH increases. Further, the DNA cleaving activity can also be detected on linear and single-stranded substrates (**Fig. 3B**). The total intensity of DNA measured in each lane was used to assess linear and single-stranded circular substrate degradation. The samples with AtYoeB were then compared to the same samples lacking the toxin to yield a percentage of remaining DNA substrate. Little variation was noted between substrates (see **Fig. S6** for additional gel images and calculations) with an average remaining DNA substrate across all those tested of 70 ± 6.8 %.

**Figure 3.**
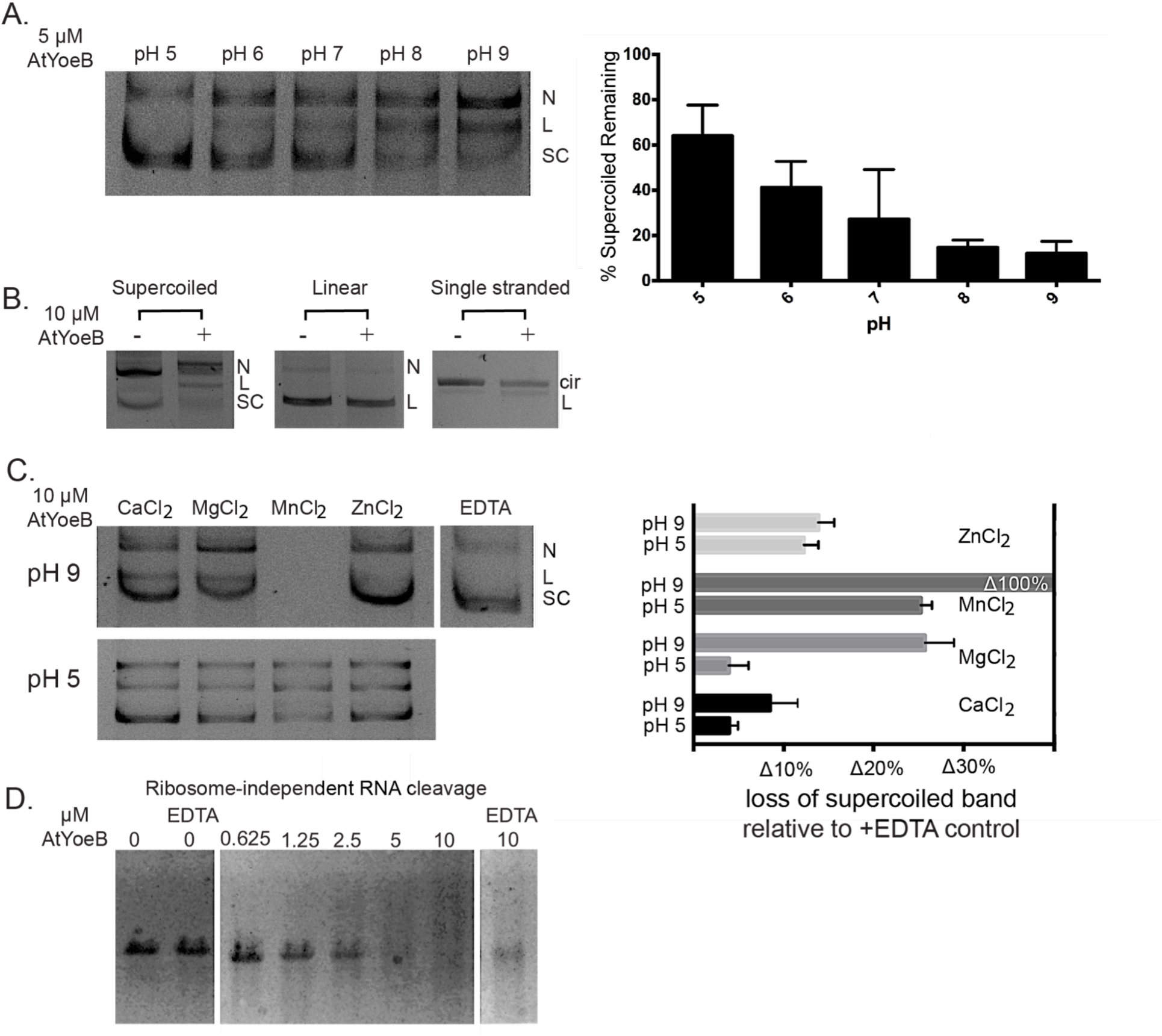
AtYoeB DNA cleavage is sensitive to pH and requires divalent cations, while RNA cleavage is independent of metal cleavage. **A.** Nuclease activity assays were carried out at the indicated pH values, and the resulting intensity of each DNA topology was quantified (*n* = 3, see Fig. S6). A marked pH dependence is noted, as pH 9 contains less of the starting supercoiled (SC) topology as it is cleaved into a nicked (N) and linear (L) topology. **B.** The DNA substrate did not affect the resulting cleavage, with approx. 25-35% decrease in total intensity for DNA after incubation with AtYoeB (*n* = 3, see Fig. S7). **C.** The DNA cleavage reaction requires a divalent cation, with manganese producing the most efficient reaction such that at pH 9 the DNA substrate is completely degraded while at pH 5 approx. 30% is degraded (*n* = 6, see Fig. S8). Magnesium is also sufficient, again with increased degradation at pH 9 versus pH 5. Zinc and calcium do not show strong pH dependence and are not as efficient at mediating catalysis. **D.** AtYoeB also degrades RNA in a dose-dependent manner in the absence of the ribosome, but this reaction does not require a divalent cation (*n* = 3). Reactions are performed with 2.5 mM MgCl_2_ except for lanes with the chelating agent EDTA. Note that 10 μM AtYoeB degrades essentially all of the RNA substrate under these conditions, and in the presence of EDTA it is still able to degrade approx. 65% of the starting substrate.

The ability of YoeB to cleave DNA was also determined to be metal-dependent, with both magnesium and manganese able to support this catalytic activity (**Fig. 3C and S7** for additional gel images and calculations). Calcium and zinc, however, were not useful for catalysis; the metal dependence was further confirmed by a lack of DNA cleavage in the presence of EDTA. The sensitivity to pH was also confirmed in the metal-dependent assay, where at the more active pH 9 the metal ion manganese catalyzes a complete loss of DNA substrate; however, at pH 5 this is slowed to result in a loss of approx. 25% of the supercoiled band (**Fig. 3C**). A similar relationship was observed with magnesium, although the extent of degradation was tempered relative to manganese, with a loss of approx. 27% at pH 9 and only approx. a 5% loss at pH 5.

Previously, the YoeB from *E. coli* was observed to cleave RNA in a ribosome-independent reaction *in vitro*.[42] In a similar assay, we have determined that the AtYoeB toxin is also able to cleave RNA *in vitro* in the absence of the ribosome (**Fig. 3D**, additional gels in **Fig. S8**). This cleavage, however, is not a metal-dependent reaction but does proceed faster with magnesium. This is consistent with proposed mechanisms that rely on the nucleophilic activity of the 2’-OH of RNA in concert with the general acid and base residues of the protein. Calcium had a weaker impact on the catalysis of RNA degradation (data not shown), while manganese and zinc caused cleavage of RNA in the absence of the AtYoeB toxin.

### AtYoeB-mediated DNA cleavage is blocked by interaction with the AtYefM antitoxin

These experiments were carried out to further demonstrate that the observed DNA cleavage arises directly from the AtYoeB toxin. The toxin and antitoxin were expressed and purified individually, and also co-expressed and purified as a complex. When assessed immediately after purification, only the AtYoeB toxin demonstrates DNA cleavage ability (**Fig. 4A**). Adding exogenous antitoxin to reactions was not able to block the DNA cleavage. Upon prolonged incubation (> 2 wks) at 4°, some minor DNase cleavage activity was noted for the co-expressed complex. We hypothesized that some degradation of antitoxin was occurring during storage, similar to what we observed for the initial inability of antitoxin to block the DNase activity. A formal experiment utilized increased temperatures to promote this presumed degradation by incubating identical aliquots of freshly purified AtYoeB-YefM complex at 4°C, 23°C, and 37°C. After one week, these samples were analyzed for DNase activity (**Fig. 4B**) and antitoxin degradation as visualized by Coomassie-stained gels and Western blots to detect the Strep-tagged toxin and His-tagged antitoxin (**Fig. 4C**). This revealed that indeed the antitoxin was becoming degraded, evidenced by multiple protein bands reacting with the anti-His antibody and an additional low molecular weight band with no reactivity on either Western blot. These experiments also verified that the toxin was unchanged. The DNase activity was prominent in the samples with antitoxin degradation, and absent in the 4° stored sample which, while having some visible antitoxin degradation, was to a lesser extent than the samples incubated at warmer temperatures.

**Figure 4.**
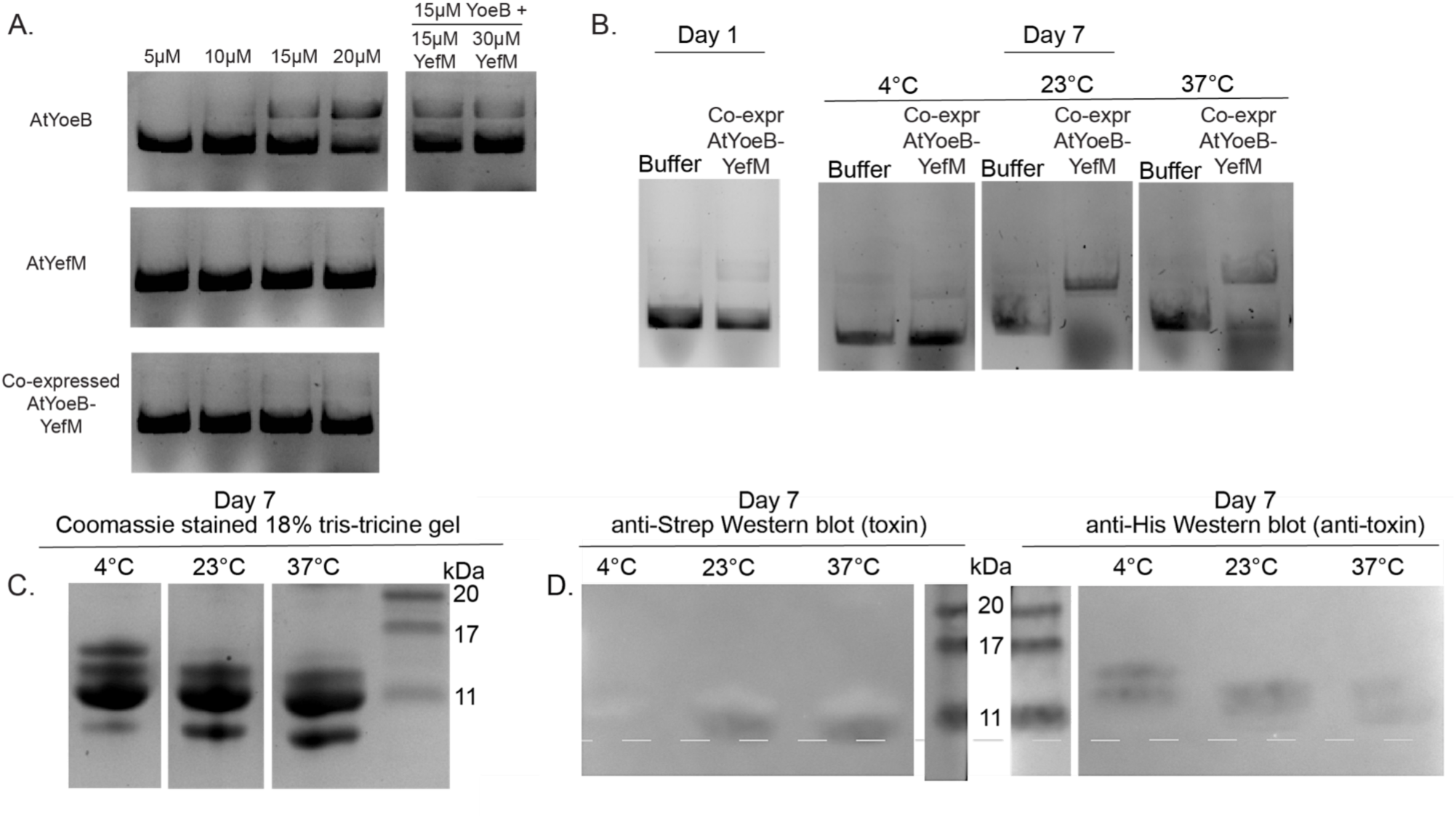
The AtYefM antitoxin is able to block the AtYoeB-mediated DNA cleavage, but it readily undergoes degradation that renders it ineffective. A. DNA degrading activity of purified toxin, antitoxin, and co-expressed complex immediately after purification. These demonstrate that only the AtYoeB toxin possesses DNA cleaving ability. Further, addition of purified antitoxin to the toxin samples is not able to block this degradation. B. DNA degradation of co-purified complex after incubation for 1 week at temperatures above 4°C results in the gain of DNase activity. C. Electrophoretic analysis of the protein samples used in panel B reveal degradation of one of the protein components even at 4°C, but with increased degradation at temperatures above 4°C. D. Western blots were used to identify the individual bands visualized in the gel in panel C, revealing that the degraded component is the His-tagged AtYefM antitoxin, while the Strep-tagged AtYoeB toxin remains unchanged.

### The dose-dependence of *in vitro* DNA and RNA cleavage are equivalent, and RNA cleavage is enhanced by the ribosome

The DNA and RNA cleavage reactions carried out by AtYoeB appear to vary in their catalytic requirements for divalent cations; however, the dose-dependence of these two cleavage reactions are equivalent (**Fig. 5A, S4, S8**). This ribosome-independent RNA cleavage has been described previously, and, additionally, it was noted that *in vivo*, the ribosome-dependent activity is the relevant reaction. We utilized an *in vitro* cell-free coupled transcription-translation reaction to assess the impact of the ribosome on the AtYoeB dose-dependent nucleic acid cleavage (**Fig. 5B**). While the output of this assay is not a gel-based measurement but is instead the production of fluorescent protein, the dose-dependence is the same regardless of if the reactions are initiated with DNA or with RNA. We were thus able to directly compare the contribution of the ribosome to catalytic efficiency, which is an improvement of approximately a factor of ten. Because the dose-dependence is the same regardless of starting with a DNA substrate, the ribosomal-dependent RNA cleavage must be the dominant reaction. This then provides the first direct measurements of equivalent nuclease activities for a toxin molecule.

**Figure 5.**
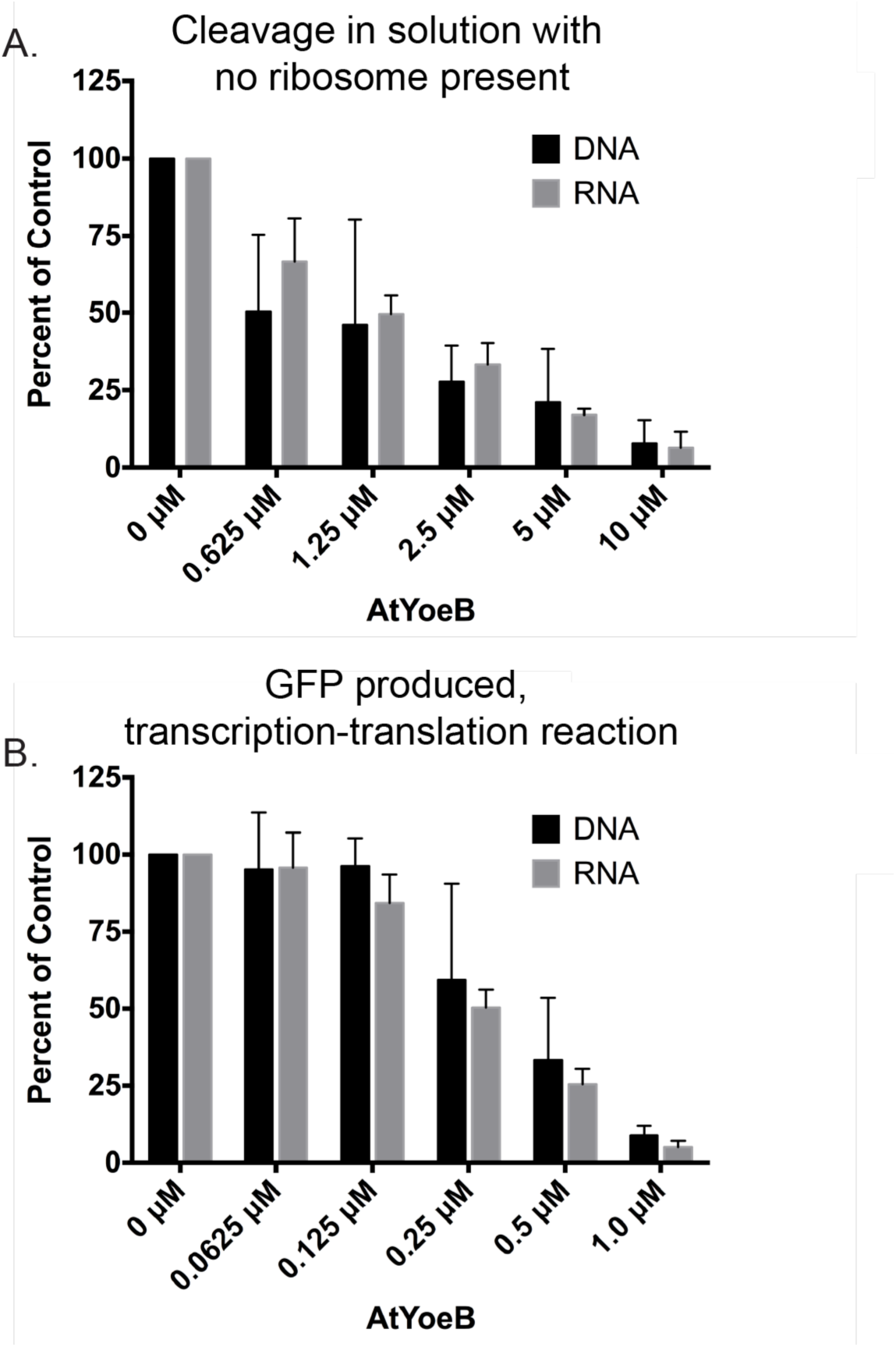
Both DNA and RNA degradation follow the same AtYoeB dose-dependence, and inclusion of the ribosome enhances cleavage by about 10x. **A.** Reactions were performed using either DNA or RNA substrates in the presence of 2.5 mM MgCl_2_ and increasing AtYoeB. The intensity of either supercoiled DNA (*n* = 3, representative shown in Fig. 3) or total RNA (*n* = 3, representative shown in Fig. 3) were measured after electrophoresis and normalized relative to the control sample with no AtYoeB. **B.** *In vitro* coupled transcription translation assays were carried out using either linearized DNA or *in vitro* transcribed RNA coding for Green Fluorescent Protein incubated with increasing concentrations of AtYoeB; the resulting fluorescence was measured and normalized to control samples containing no AtYoeB (*n* = 3).

## SUMMARY AND CONCLUSIONS

As opposed to previously studied YoeB toxins, AtYoeB readily produces soluble toxin in the absence of antitoxin, providing an ideal system to further explore YoeB activity. While its structure and sequence are highly similar to the well-characterized YoeB toxin from *E. coli*,. [39-44] specific sequence changes can be pinpointed at toxin-antitoxin interaction points. This would limit cross-reactivity of the antitoxins from different YoeB-YefM operons. Continued experiments are seeking to understand the basis for the loss of AtYoeB toxicity with the hypothesis that the interactions with the *Agrobacterium tumefaciens* ribosome must be distinct. However, no significant sequence changes at the ribosome-interacting regions are noted.

The AtYoeB-YefM complex is a heterotetramer. Individual toxin and antitoxin are dimeric, and they interact with canonically tight affinity in the high picomolar range. This high affinity can also be inferred by an increased melting transition point of approx. 20 °C, while the AtYefM antitoxin does not contain sufficient folded structure to yield a hydrophobic core as needed to produce a signal in this assay.

It is known that certain toxins in the Rel-superfamily such as MazF, YafQ, and RelE cleave truncated mRNA at the ribosomal A-site; however, MazF has also been reported to function independently of the ribosome.[61] Briefly, the toxin will bind to a ribosomal subunit, interacting with the A site and preventing formation of the translation initiation complex.[19, 42-44, 65] Additionally, this allows the cleavage of translated mRNA at the A site, releasing the 3’-end of the mRNA from the ribosome.[43] Similarly, we find that the YoeB toxin from *A. tumefaciens* is able to cleave RNA in both ribosome-dependent and –independent reactions.

These experiments demonstrate that the antitoxin protein is prone to degradation, such that when purified in the absence of toxin it is then unable to form interactions with the AtYoeB toxin that block DNase activity, even at a two times molar excess. This is likely due to an increased rate in the degradation of YefM upon purification in the absence of the toxin protein. The co-expressed complex of AtYoeB and AtYefM clearly form a productive complex that is devoid of DNase activity; the gain of this function, then, correlated with the visible degradation of antitoxin confirms that the AtYoeB toxin is capable of direct DNA degradation, at least *in vitro*.

AtYoeB, and likely other YoeB toxins, possess the necessary residues to act as a general nuclease, cleaving both RNA and DNA. While this activity is seen in the presence and absence of the ribosome, the presence of the ribosome appears to order the C-terminal catalytic residue, yielding an activity 10 times that observed in ribosome-independent reactions. The presence of magnesium in solution can somewhat mimic binding of the ribosome, increasing the rate of RNA cleavage. In the absence of the 2’-OH group, Mg2+ is essential for DNA cleavage – a potential mechanism based on DNaseI and related HNH nucleases that utilize two histidine residues as opposed to the histidine and glutamic acid that cleave RNA.[66] Due to the involvement of two histidine residues in the cleavage of DNA, this reaction would be sensitive to pH changes. Coupled with the partial charge distribution on the phosphoryl oxygens due to magnesium coordination, these different local pKa environments of the histidine residues could favor catalysis.

It is striking that both RNA nuclease and DNA nicking activity has been identified for the VapD toxin family.[67] The active site residues are shared by both nuclease activities, with acidic residues serving as the catalytic residues that coordinate a metal binding site. This mechanism was noted to be pH dependent, with increased activity above pH 8, similar to what is described in the current study. However, analysis of the structures of VapD and YoeB reveal no structural homology. Other studies have noted the conservation of the RelE/ParE fold, including homology with other RNase proteins including colicins and barnase.[68] The recognition of this deoxyribonuclease activity as a similar and potentially overlapping mechanism with the RNase ability highlights the plasticity of the conserved fold found in YoeB toxins.

## ASSOCIATED CONTENT

### Author Contributions

This study was conceptualized by JRA and CRB. All authors contributed to data collection; specifically, TM collected all the RNase data and, JRA, TM and JM collected the DNase data;

JM developed the experiments and collected the data in Fig 4; ER carried out the crystallographic study; CRB carried out the DSF and MALS experiments, and JRA carried out the BLI experiments. Data analysis was performed by JRA, TM, and CRB. The manuscript was written by JRA and CRB, and all authors edited and approved the final version.

### Funding Sources

Research reported in this publication was supported by an Institutional Development Award (IDeA) from the National Institute of General Medical Sciences of the National Institutes of Health under grant number P20GM103640.

The authors declare no competing financial interests.

### Accession Codes

*Atu2017*, UniProt ID A9CID9; *Atu2018*, UniProt ID Q7CY23; AtYoeB, PDB ID 6N90

## ACKNOWLEDGMENT

The authors are grateful to Dr. Ben F. Holt III for kindly providing the *Agrobacterium tumefaciens* genomic DNA, the University of Oklahoma Protein Production Core Facility for assistance with sample preparation, and the University of Oklahoma Macromolecular Crystallography Laboratory for assistance with crystallization and X-ray data collection; both of these facilities are supported by an Institutional Development Award (IDeA) from the National Institute of General Medical Sciences of the National Institutes of Health under grant number P20GM103640. We would also like to thank Hannah Panlilio for collection of initial MALS data, John C. White for pilot studies, and Dr. Meenakumari Muthuramalingham for cloning. We are grateful to Dr. Paul Sims (University of Oklahoma) and to the Scanlan group (Trinity College, Dublin) for discussions on DNase catalytic mechanisms. Data collection for the AtYoeB crystal structure was carried out at the Stanford Synchrotron Radiation Lightsource, SLAC National Accelerator Laboratory, which is supported by the U.S. Department of Energy, Office of Science, Office of Basic Energy Sciences under Contract No. DE-AC02-76SF00515. The SSRL Structural Molecular Biology Program is supported by the DOE Office of Biological and Environmental Research, and by the National Institutes of Health, National Institute of General Medical Sciences (including P41GM103393).

## Supplemental Materials

**Table S1.**
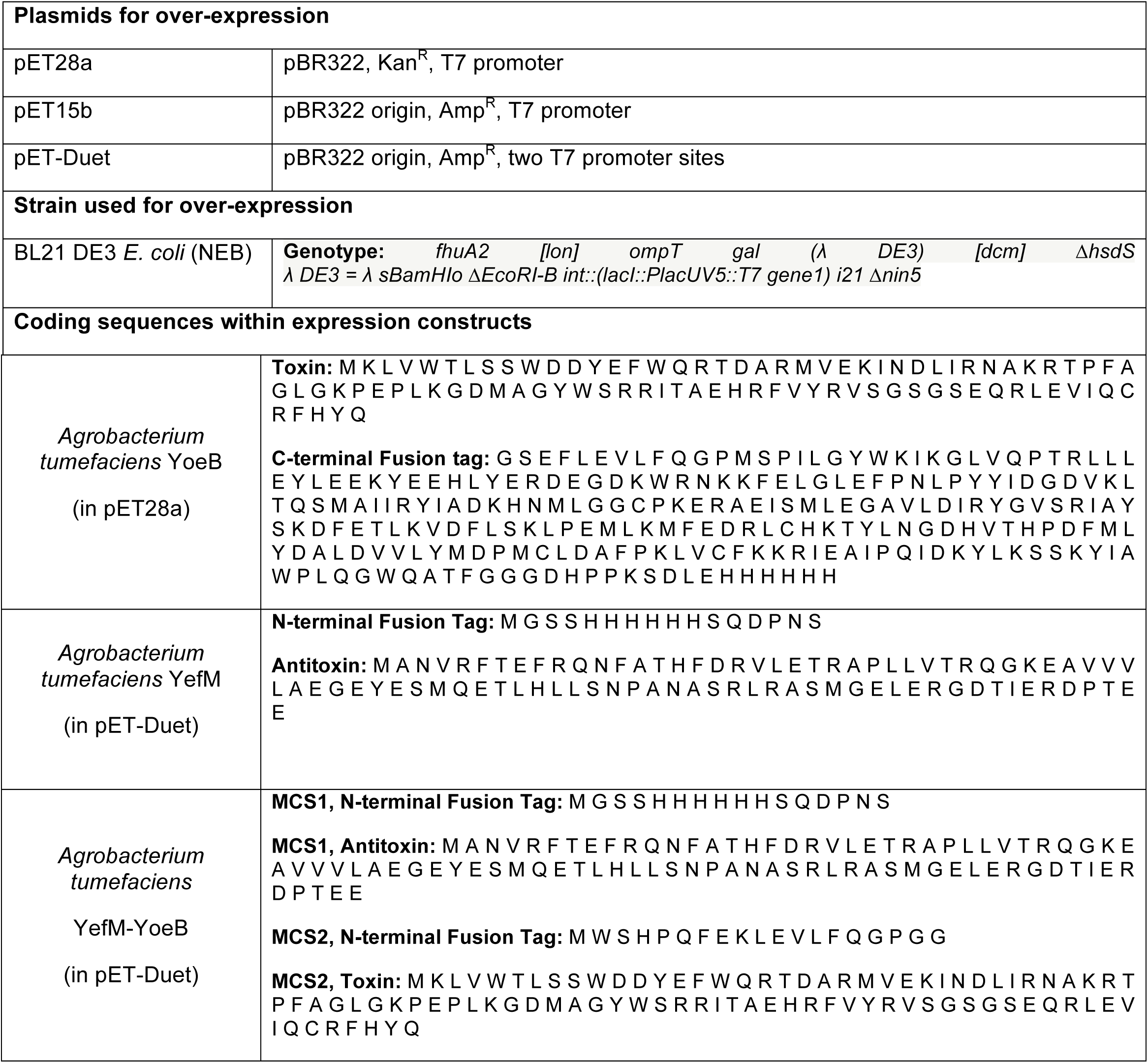
Materials used in these studies.

**Figure S1.**
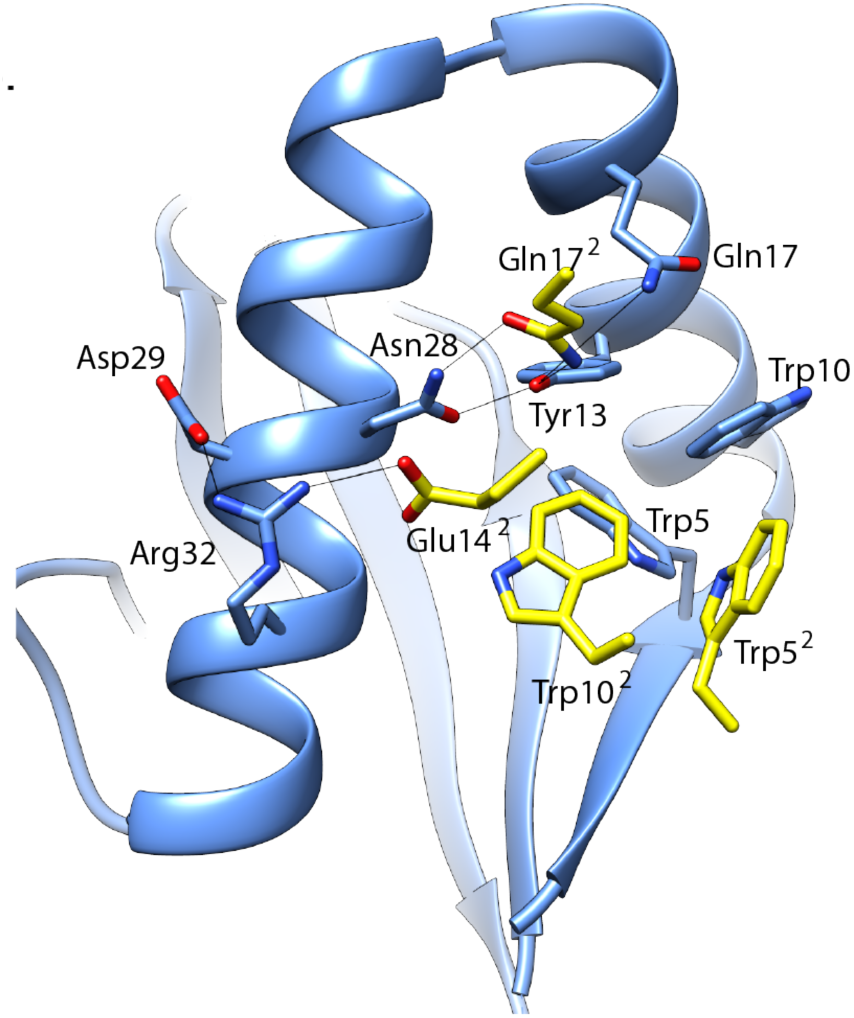
The AtYoeB dimer interface has three amino acid differences as compared to the EcYoeB. The dimer interface of AtYoeB contains the canonical Trp stack, comprised of Trp5 and Trp10, in addition to numerous stabilizing polar interactions. The identity at position 14 in AtYoeB is polar and contributes additional electrostatic interactions as compared to the EcYoeB. Changes at position 18 and 32 are more conservative and maintain the electrostatic interactions. Note that the second molecule is obscured for a clearer view; further, only one set of the symmetrical interactions are shown.

**Figure S2.**
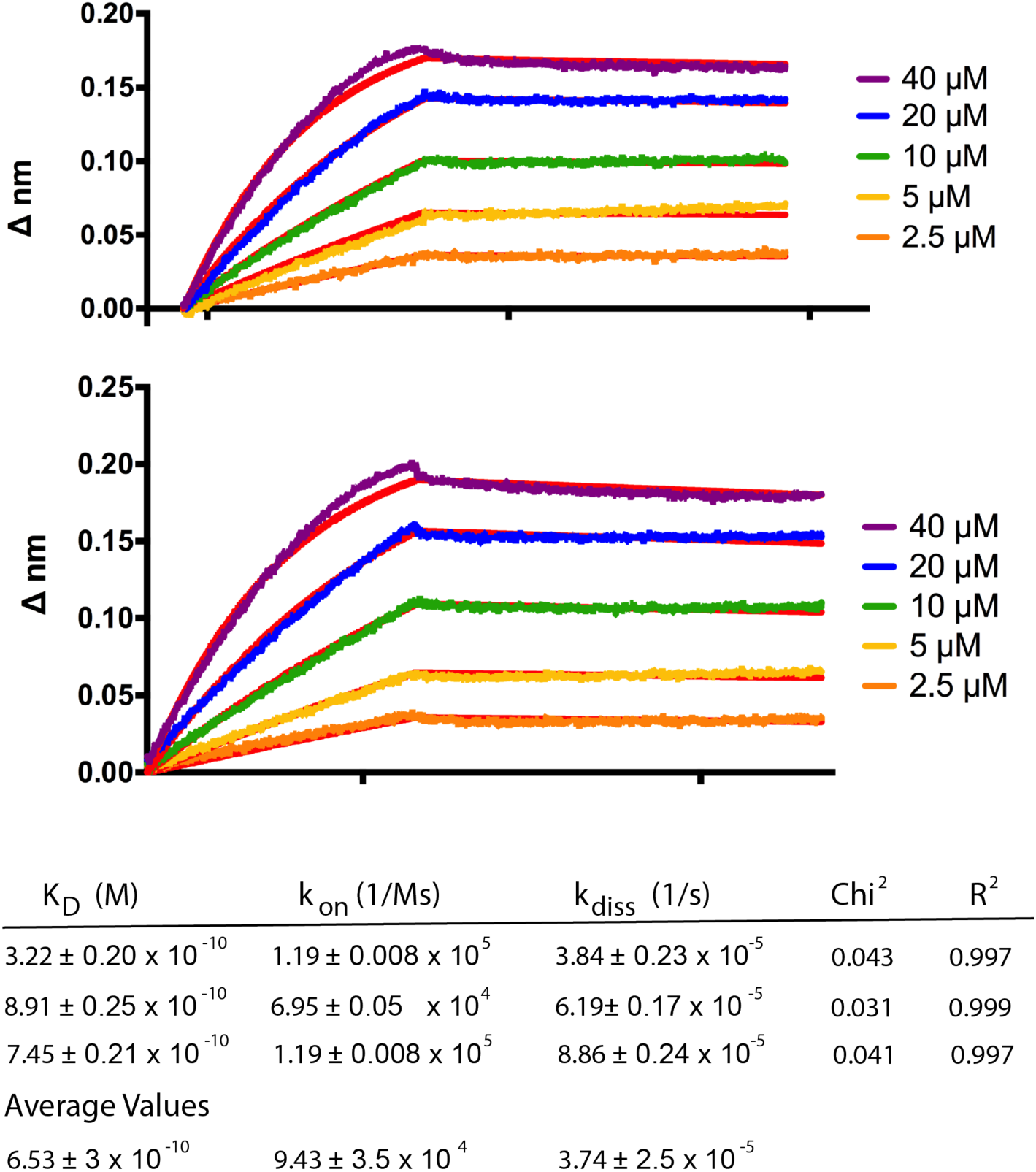
BLI data used to calculate an interaction strength between AtYoeB and AtYefM of 653 pM.

**Figure S3.**
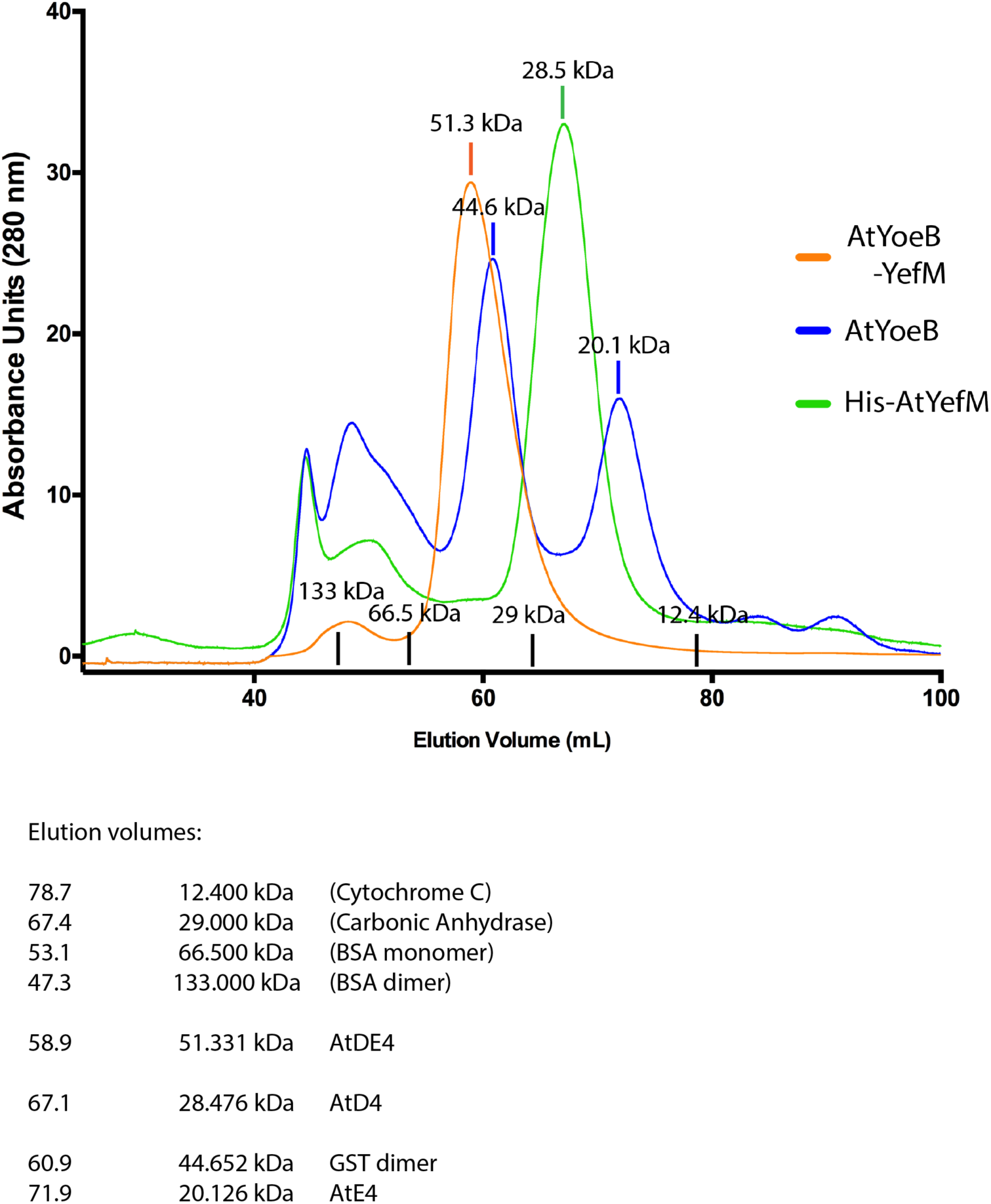
Size exclusion analysis of AtYoeB after cleavage of its GST-His fusion affinity tag, of AtYefM with its His affinity tag, and of the co-expressed AtYoeB-YefM complex (containing a His tag on the YefM and a Strep tag on the YoeB). Each sample was first purified by Ni-NTA, concentrated, and directly applied to a Sephadex S-75 column. Note that the co-expressed sample does not contain additional peaks that would correspond to any other oligomeric species aside from the heterotetramer. Standard sized proteins used to calibrate the column are listed, as are the calculated molecular masses based on elution position.

**Figure S4.**
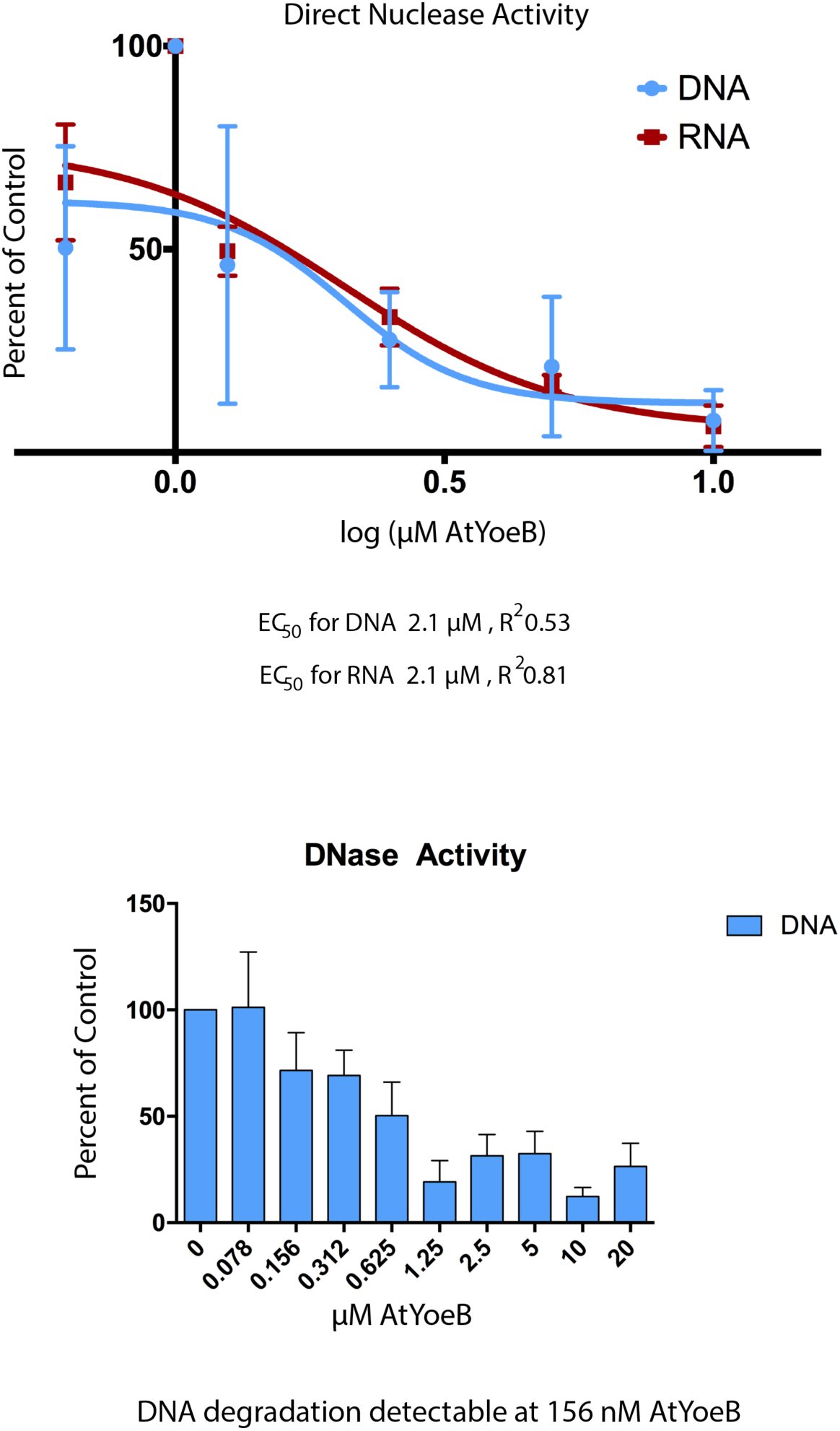
Calculation of the EC_50_ value for DNase and RNase activity *in vitro* and in the absence of the ribosome. DNase activity is detected at concentrations as low as 156 nM.

**Figure S5.**
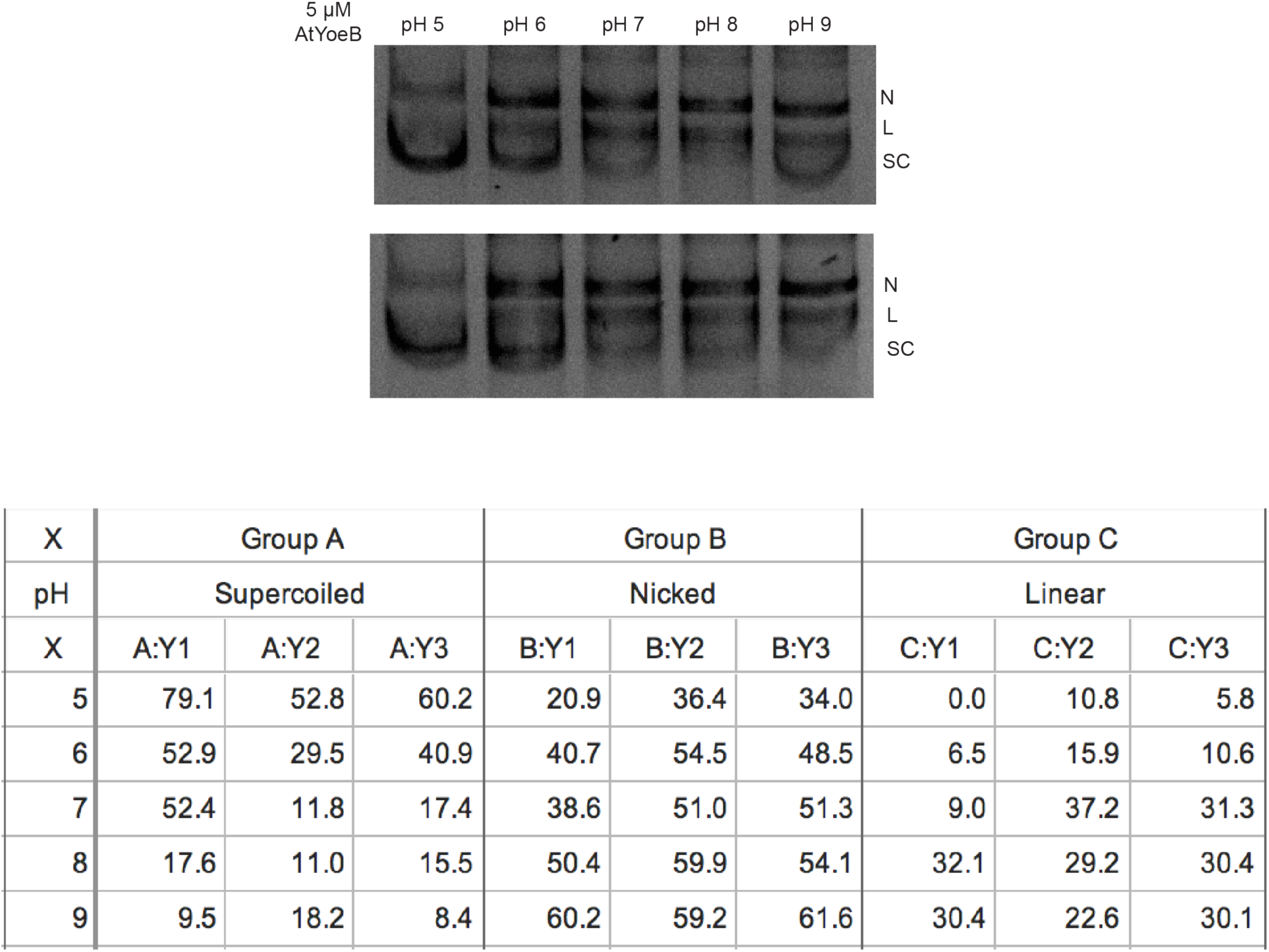
Replicates and calculations for the pH dependence of the observed AtYoeB-mediated DNase activity.

**Figure S6.**
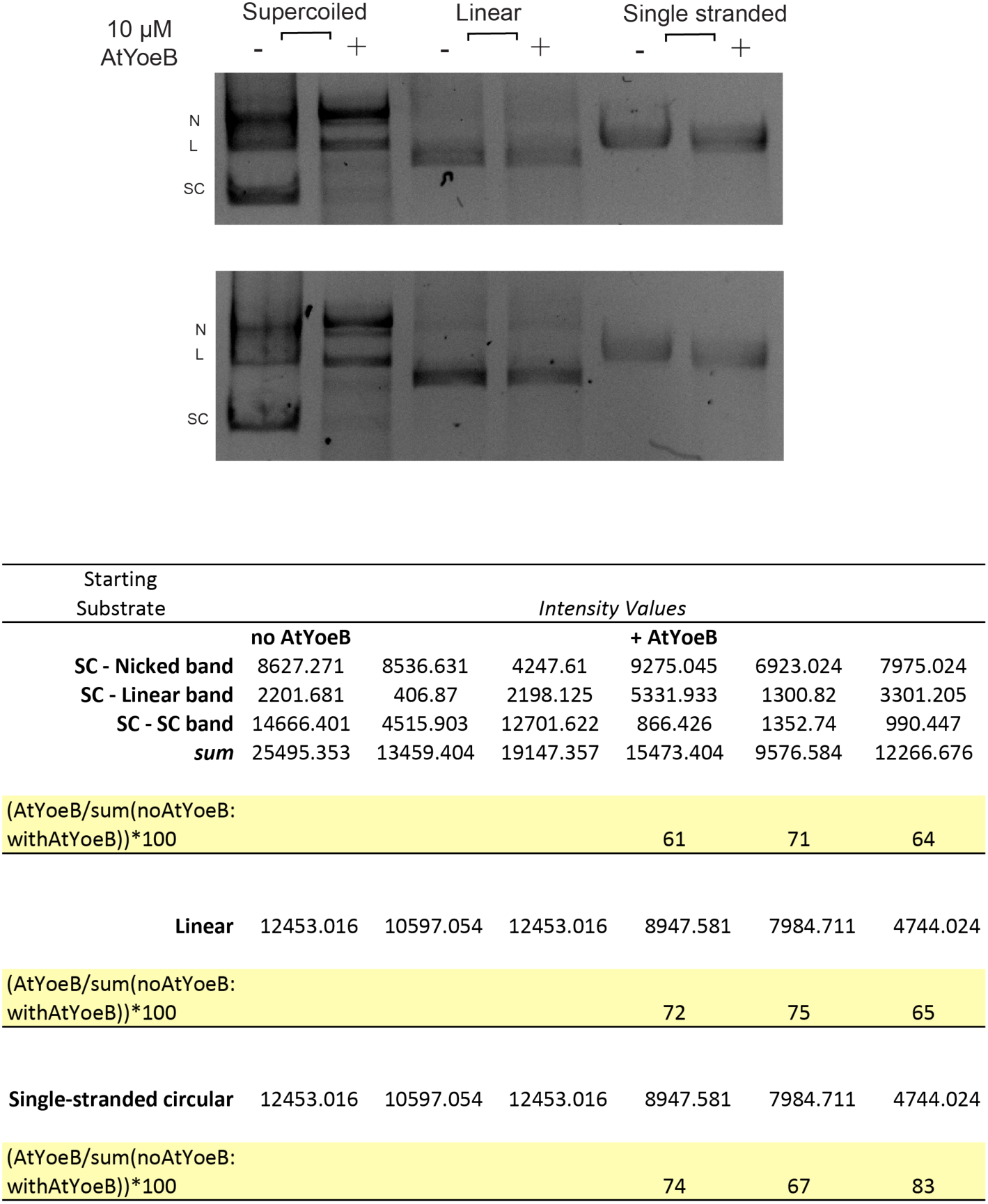
Replicates and calculations for the substrate independence of the observed AtYoeB-mediated DNase activity.

**Figure S7.**
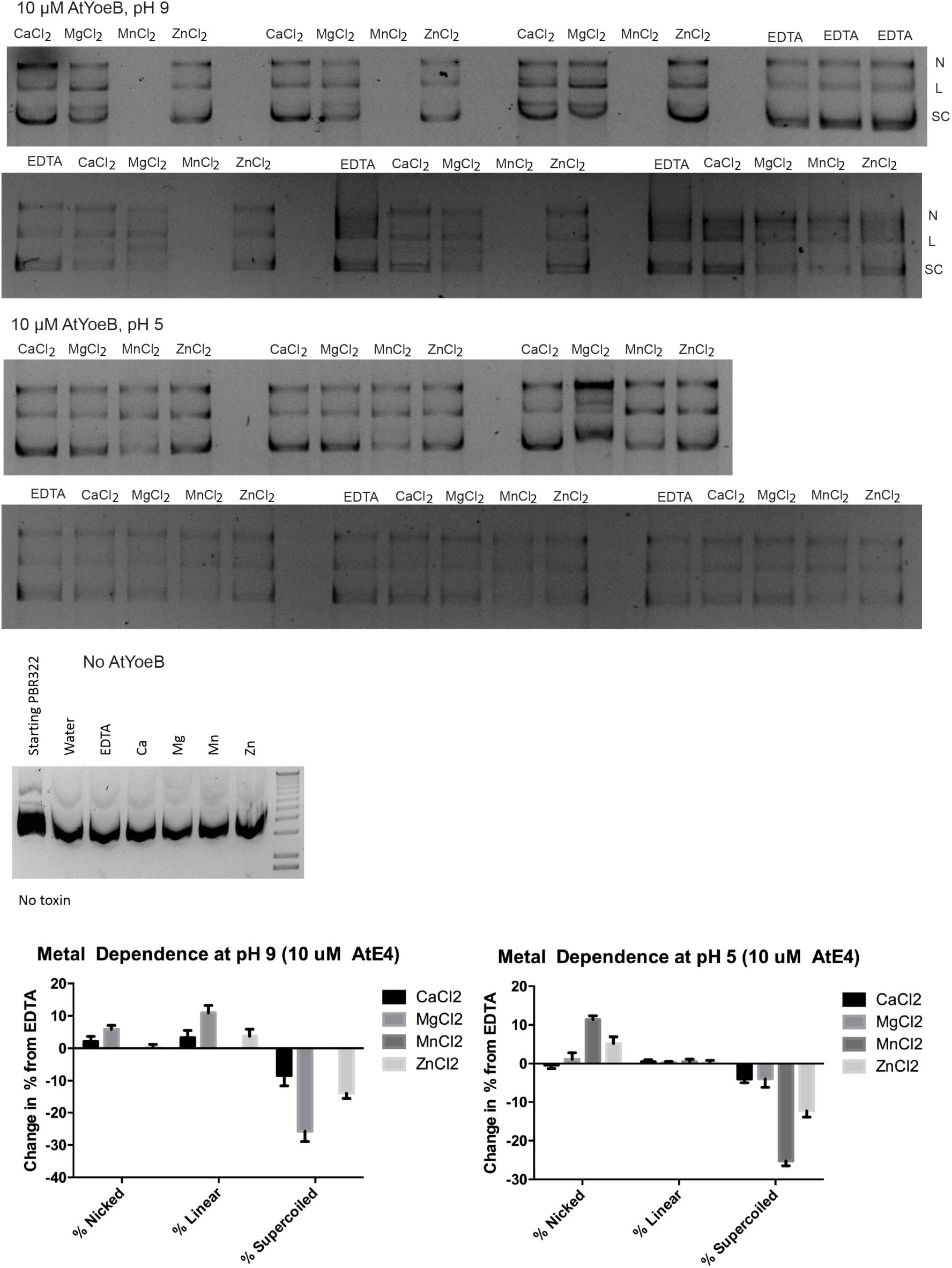
Replicates and calculations for the divalent cation dependence of the observed AtYoeB-mediated DNase activity.

**Figure S8.**
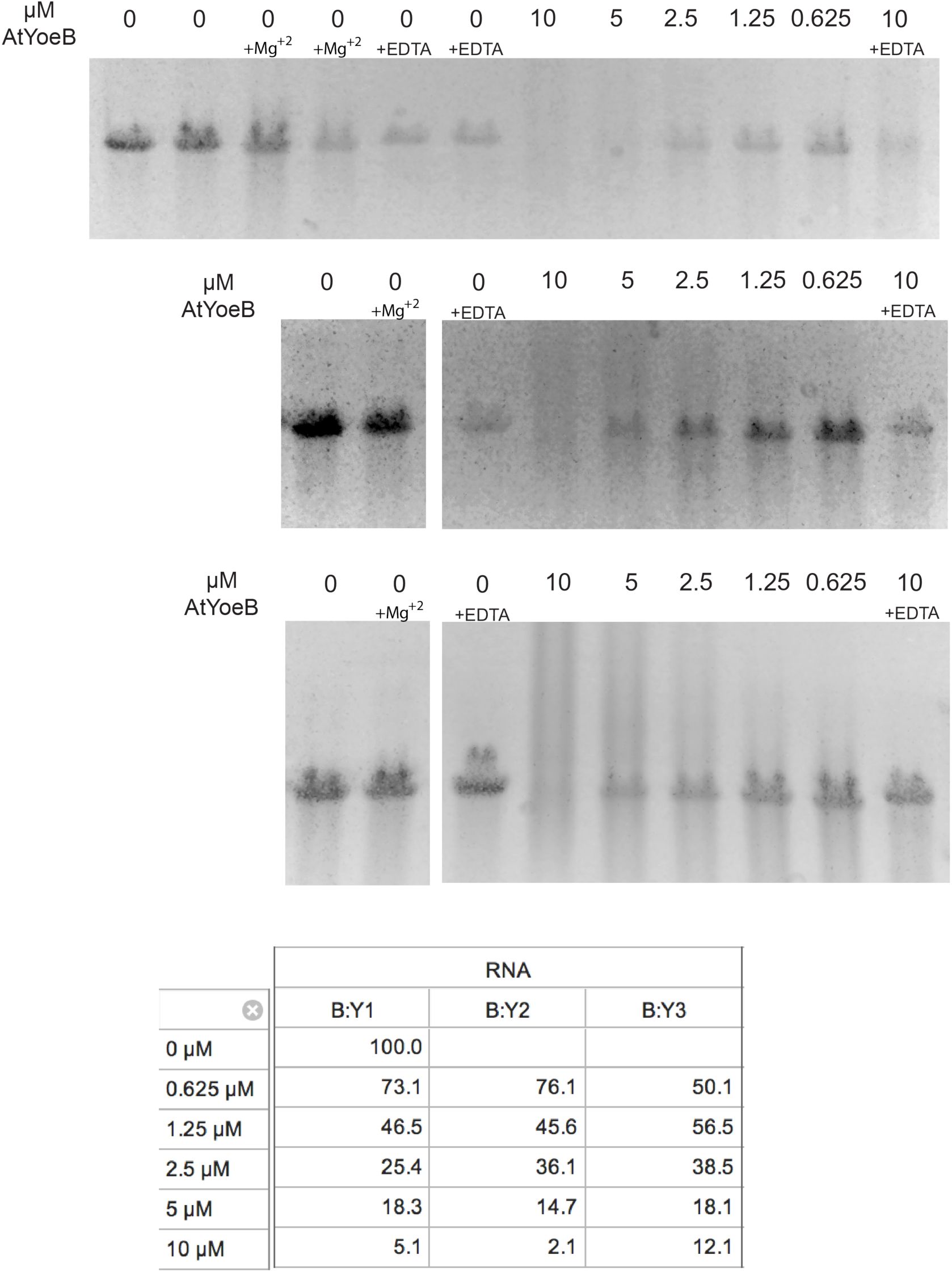
Replicates for the dose dependence of the observed AtYoeB-mediated RNase activity in the absence of the ribosome, also demonstrating a lack of metal dependence. Reactions utilized 80 ng of RNA incubated with increasing concentrations of AtYoeB, and non-specific degradation was visualized by measuring the loss of intensity of the RNA as compared to the control. Magnesium was included as noted at 2.5 mM, while EDTA was present where indicated at 5 mM.

## REFERENCES

1. Pandey DP, Gerdes K: Toxin-antitoxin loci are highly abundant in free-living but lost from host-associated prokaryotes. Nucleic Acids Res 2005, 33(3):966–976.

2. Unterholzner SJ, Poppenberger B, Rozhon W: Toxin-antitoxin systems: Biology, identification, and application. Mobile genetic elements 2013, 3(5):e26219.

3. Hall AM, Gollan B, Helaine S: Toxin-antitoxin systems: reversible toxicity. Current opinion in microbiology 2017, 36:102-110.

4. Harms A, Brodersen DE, Mitarai N, Gerdes K: Toxins, Targets, and Triggers: An Overview of Toxin-Antitoxin Biology. Molecular cell 2018, 70(5):768–784.

5. Ronneau S, Helaine S: Clarifying the Link between Toxin-Antitoxin Modules and Bacterial Persistence. Journal of molecular biology 2019.

6. Hayes F, Kedzierska B: Regulating toxin-antitoxin expression: controlled detonation of intracellular molecular timebombs. Toxins 2014, 6(1):337–358.

7. Muthuramalingam M, White JC, Bourne CR: Toxin-antitoxin modules are pliable switches activated by multiple protease pathways. Toxins 2016, 8(7):214–229.

8. Brzozowska I, Zielenkiewicz U: Regulation of toxin-antitoxin systems by proteolysis. Plasmid 2013, 70(1):33–41.

9. Lehnherr H, Yarmolinsky MB: Addiction protein Phd of plasmid prophage P1 is a substrate of the ClpXP serine protease of *Escherichia coli*. Proceedings of the National Academy of Sciences of the United States of America 1995, 92(8):3274–3277.

10. Page R, Peti W: Toxin-antitoxin systems in bacterial growth arrest and persistence. Nature chemical biology 2016, 12(4):208–214.

11. Tsuchimoto S, Ohtsubo E: Autoregulation by cooperative binding of the PemI and PemK proteins to the promoter region of the pem operon. Mol Gen Genet 1993, 237(1-2):81-88.

12. Van Melderen L, Bernard P, Couturier M: Lon-dependent proteolysis of CcdA is the key control for activation of CcdB in plasmid-free segregant bacteria. Molecular microbiology 1994, 11(6):1151–1157.

13. Donegan NP, Thompson ET, Fu Z, Cheung AL: Proteolytic regulation of toxin-antitoxin systems by ClpPC in *Staphylococcus aureus*. Journal of bacteriology 2010, 192(5):1416–1422.

14. Lehnherr H, Maguin E, Jafri S, Yarmolinsky MB: Plasmid addiction genes of bacteriophage P1: doc, which causes cell death on curing of prophage, and phd, which prevents host death when prophage is retained. Journal of molecular biology 1993, 233(3):414–428.

15. Gerdes K, Rasmussen PB, Molin S: Unique type of plasmid maintenance function: postsegregational killing of plasmid-free cells. Proceedings of the National Academy of Sciences of the United States of America 1986, 83(10):3116–3120.

16. Ogura T, Hiraga S: Mini-F plasmid genes that couple host cell division to plasmid proliferation. Proceedings of the National Academy of Sciences of the United States of America 1983, 80(15):4784–4788.

17. Kedzierska B, Hayes F: Emerging Roles of Toxin-Antitoxin Modules in Bacterial Pathogenesis. Molecules 2016, 21(6).

18. Winther KS, Gerdes K: Enteric virulence associated protein VapC inhibits translation by cleavage of initiator tRNA. Proceedings of the National Academy of Sciences of the United States of America 2011, 108(18):7403–7407.

19. Christensen SK, Gerdes K: RelE toxins from bacteria and Archaea cleave mRNAs on translating ribosomes, which are rescued by tmRNA. Molecular microbiology 2003, 48(5):1389–1400.

20. Pedersen K, Zavialov AV, Pavlov MY, Elf J, Gerdes K, Ehrenberg M: The bacterial toxin RelE displays codon-specific cleavage of mRNAs in the ribosomal A site. Cell 2003, 112(1):131–140.

21. Castro-Roa D, Garcia-Pino A, De Gieter S, van Nuland NAJ, Loris R, Zenkin N: The Fic protein Doc uses an inverted substrate to phosphorylate and inactivate EF-Tu. Nat Chem Biol 2013, 9(12):811–817.

22. Jorgensen MG, Pandey DP, Jaskolska M, Gerdes K: HicA of *Escherichia coli* defines a novel family of translation-independent mRNA interferases in bacteria and archaea. Journal of bacteriology 2009, 191(4):1191–1199.

23. Kawano M, Aravind L, Storz G: An antisense RNA controls synthesis of an SOS-induced toxin evolved from an antitoxin. Molecular microbiology 2007, 64(3):738–754.

24. Zhang Y, Zhang J, Hoeflich KP, Ikura M, Qing G, Inouye M: MazF cleaves cellular mRNAs specifically at ACA to block protein synthesis in Escherichia coli. Molecular cell 2003, 12(4):913–923.

25. Jiang Y, Pogliano J, Helinski DR, Konieczny I: ParE toxin encoded by the broad-host-range plasmid RK2 is an inhibitor of *Escherichia coli* gyrase. Molecular microbiology 2002, 44(4):971–979.

26. Bernard P, Couturier M: Cell killing by the F plasmid CcdB protein involves poisoning of DNA-topoisomerase II complexes. Journal of molecular biology 1992, 226(3):735–745.

27. Harms A, Stanger FV, Scheu PD, de Jong IG, Goepfert A, Glatter T, Gerdes K, Schirmer T, Dehio C: Adenylylation of Gyrase and Topo IV by FicT Toxins Disrupts Bacterial DNA Topology. Cell Rep 2015, 12(9):1497–1507.

28. Grady R, Hayes F: Axe-Txe, a broad-spectrum proteic toxin-antitoxin system specified by a multidrug-resistant, clinical isolate of *Enterococcus faecium*. Molecular microbiology 2003, 47(5):1419–1432.

29. Guglielmini J, Van Melderen L: Bacterial toxin-antitoxin systems: Translation inhibitors everywhere. Mobile genetic elements 2011, 1(4):283–290.

30. Leplae R, Geeraerts D, Hallez R, Guglielmini J, Dreze P, Van Melderen L: Diversity of bacterial type II toxin-antitoxin systems: a comprehensive search and functional analysis of novel families. Nucleic acids research 2011, 39(13):5513–5525.

31. Park SJ, Son WS, Lee BJ: Structural overview of toxin-antitoxin systems in infectious bacteria: A target for developing antimicrobial agents. Biochim Biophys Acta 2013, 1843(6):1155–1167.

32. Schmidt O, Schuenemann VJ, Hand NJ, Silhavy TJ, Martin J, Lupas AN, Djuranovic S: prlF and yhaV encode a new toxin-antitoxin system in Escherichia coli. Journal of molecular biology 2007, 372(4):894–905.

33. Polom D, Boss L, Wegrzyn G, Hayes F, Kedzierska B: Amino acid residues crucial for specificity of toxin-antitoxin interactions in the homologous Axe-Txe and YefM-YoeB complexes. The FEBS journal 2013, 280(22):5906–5918.

34. Dy RL, Richter C, Salmond GP, Fineran PC: Remarkable Mechanisms in Microbes to Resist Phage Infections. Annu Rev Virol 2014, 1(1):307–331.

35. Magnuson RD: Hypothetical functions of toxin-antitoxin systems. Journal of bacteriology 2007, 189(17):6089–6092.

36. Van Melderen L: Toxin-antitoxin systems: why so many, what for? Current opinion in microbiology 2010, 13(6):781–785.

37. Van Melderen L, Saavedra De Bast M: Bacterial toxin-antitoxin systems: more than selfish entities? PLoS Genet 2009, 5(3):e1000437.

38. Muthuramalingam M, White JC, Murphy T, Ames JR, Bourne CR: The toxin from a ParDE toxin-antitoxin system found in *Pseudomonas aeruginosa* offers protection to cells challenged with anti-gyrase antibiotics. Molecular microbiology 2018.

39. Chan WT, Nieto C, Harikrishna JA, Khoo SK, Othman RY, Espinosa M, Yeo CC: Genetic regulation of the yefM-yoeB toxin-antitoxin locus of Streptococcus pneumoniae. Journal of bacteriology 2011, 193(18):4612–4625.

40. Cherny I, Rockah L, Gazit E: The YoeB toxin is a folded protein that forms a physical complex with the unfolded YefM antitoxin. Implications for a structural-based differential stability of toxin-antitoxin systems. The Journal of biological chemistry 2005, 280(34):30063–30072.

41. Feng S, Chen Y, Kamada K, Wang H, Tang K, Wang M, Gao YG: YoeB-ribosome structure: a canonical RNase that requires the ribosome for its specific activity. Nucleic acids research 2013, 41(20):9549–9556.

42. Kamada K, Hanaoka F: Conformational change in the catalytic site of the ribonuclease YoeB toxin by YefM antitoxin. Molecular cell 2005, 19(4):497–509.

43. Pavelich IJ, Maehigashi T, Hoffer ED, Ruangprasert A, Miles SJ, Dunham CM: Monomeric YoeB toxin retains RNase activity but adopts an obligate dimeric form for thermal stability. Nucleic acids research 2019.

44. Zhang Y, Inouye M: The inhibitory mechanism of protein synthesis by YoeB, an *Escherichia coli* toxin. The Journal of biological chemistry 2009, 284(11):6627–6638.

45. Diaz-Orejas R, Espinosa M, Yeo CC: The Importance of the Expendable: Toxin-Antitoxin Genes in Plasmids and Chromosomes. Frontiers in microbiology 2017, 8:1479.

46. Wozniak RA, Waldor MK: A toxin-antitoxin system promotes the maintenance of an integrative conjugative element. PLoS genetics 2009, 5(3):e1000439.

47. Yamamoto S, Kiyokawa K, Tanaka K, Moriguchi K, Suzuki K: Novel toxin-antitoxin system composed of serine protease and AAA-ATPase homologues determines the high level of stability and incompatibility of the tumor-inducing plasmid pTiC58. Journal of bacteriology 2009, 191(14):4656–4666.

48. Davis TL, Helinski DR, Roberts RC: Transcription and autoregulation of the stabilizing functions of broad-host-range plasmid RK2 in Escherichia coli, Agrobacterium tumefaciens and Pseudomonas aeruginosa. Molecular microbiology 1992, 6(14):1981–1994.

49. Robert X, Gouet P: Deciphering key features in protein structures with the new ENDscript server. Nucleic acids research 2014, 42(Web Server issue):W320–324.

50. Pettersen EF, Goddard TD, Huang CC, Couch GS, Greenblatt DM, Meng EC, Ferrin TE: UCSF Chimera--a visualization system for exploratory research and analysis. Journal of Computational Chemistry 2004, 25(13):1605–1612.

51. Kabsch W: Xds. Acta crystallographica Section D, Biological crystallography 2010, 66(Pt 2):125-132.

52. Evans P: Scaling and assessment of data quality. Acta crystallographica Section D, Biological crystallography 2006, 62(Pt 1):72-82.

53. Collaborative Computational Project N: The CCP4 suite: programs for protein crystallography. Acta crystallographica Section D, Biological crystallography 1994, 50(Pt 5):760–763.

54. McCoy AJ, Grosse-Kunstleve RW, Adams PD, Winn MD, Storoni LC, Read RJ: Phaser crystallographic software. J Appl Crystallogr 2007, 40(Pt 4):658–674.

55. Emsley P, Lohkamp B, Scott WG, Cowtan K: Features and development of Coot. Acta crystallographica 2010, 66(Pt 4):486–501.

56. Afonine PV, Grosse-Kunstleve RW, Echols N, Headd JJ, Moriarty NW, Mustyakimov M, Terwilliger TC, Urzhumtsev A, Zwart PH, Adams PD: Towards automated crystallographic structure refinement with phenix.refine. Acta crystallographica Section D, Biological crystallography 2012, 68(Pt 4):352–367.

57. Schindelin J, Rueden CT, Hiner MC, Eliceiri KW: The ImageJ ecosystem: An open platform for biomedical image analysis. Molecular Reproduction and Development 2015, 82(7-8):518-529.

58. Sevin EW, Barloy-Hubler F: RASTA-Bacteria: a web-based tool for identifying toxin-antitoxin loci in prokaryotes. Genome Biol 2007, 8(8):R155.

59. Xie Y, Wei Y, Shen Y, Li X, Zhou H, Tai C, Deng Z, Ou HY: TADB 2.0: an updated database of bacterial type II toxin–antitoxin loci. Nucleic acids research 2018, 46(D1):D749-D753.

60. Arbing MA, Handelman SK, Kuzin AP, Verdon G, Wang C, Su M, Rothenbacher FP, Abashidze M, Liu M, Hurley JM et al: Crystal structures of Phd-Doc, HigA, and YeeU establish multiple evolutionary links between microbial growth-regulating toxin-antitoxin systems. Structure 2010, 18(8):996–1010.

61. Maehigashi T, Ruangprasert A, Miles SJ, Dunham CM: Molecular basis of ribosome recognition and mRNA hydrolysis by the *E. coli* YafQ toxin. Nucleic acids research 2015, 43(16):8002–8012.

62. Aakre CD, Herrou J, Phung TN, Perchuk BS, Crosson S, Laub MT: Evolving new protein-protein interaction specificity through promiscuous intermediates. Cell 2015, 163(3):594–606.

63. Ames JR, Muthuramalingam M, Murphy T, Najar FZ, Bourne CR: Expression of different ParE toxins results in conserved phenotypes with distinguishable classes of toxicity. Microbiologyopen 2019:e902.

64. Kumar P, Issac B, Dodson EJ, Turkenburg JP, Mande SC: Crystal structure of Mycobacterium tuberculosis YefM antitoxin reveals that it is not an intrinsically unstructured protein. Journal of molecular biology 2008, 383(3):482–493.

65. Christensen SK, Maenhaut-Michel G, Mine N, Gottesman S, Gerdes K, Van Melderen L: Overproduction of the Lon protease triggers inhibition of translation in *Escherichia coli:* involvement of the yefM-yoeB toxin-antitoxin system. Molecular microbiology 2004, 51(6):1705–1717.

66. Suck D, Oefner C: Structure of DNase I at 2.0 A resolution suggests a mechanism for binding to and cutting DNA. Nature 1986, 321(6070):620–625.

67. Lee KY, Lee KY, Kim JH, Lee IG, Lee SH, Sim DW, Won HS, Lee BJ: Structure-based functional identification of Helicobacter pylori HP0268 as a nuclease with both DNA nicking and RNase activities. Nucleic acids research 2015, 43(10):5194–5207.

68. Gucinski GC, Michalska K, Garza-Sanchez F, Eschenfeldt WH, Stols L, Nguyen JY, Goulding CW, Joachimiak A, Hayes CS: Convergent Evolution of the Barnase/EndoU/Colicin/RelE (BECR) Fold in Antibacterial tRNase Toxins. Structure 2019.

